# CGAgentX: Agentic AI Framework to Develop Transferable Coarse-Grained Models

**DOI:** 10.64898/2026.04.17.719081

**Authors:** Swarnadeep Seth, Sanket A. Deshmukh

## Abstract

We present CGAgentX, a general autonomous multi-agent framework in which specialized LLM-based agents coordinate the optimization of coarse-grained (CG) model parameters to reproduce target properties. Using polar solvents — dimethyl sulfoxide (DMSO) and N,N-dimethylacetamide (DMA) — as representative case studies, we demonstrate the framework’s capability to develop CG models that accurately reproduce key properties from atomistic simulations and experimental literature. Six specialized agents — Mapping, Topology, Boundary, Hypothesis, Diagnostic, and Optimization — operate under a Master Agent that orchestrates closed-loop, iterative parameter refinement by autonomously invoking external tools, including molecular dynamics (MD) simulations and analysis workflows, and evaluating outputs through a fitness function. Central to the framework is a Hypothesis Agent that generates and verifies physically motivated parameter hypotheses by coordinating parallel multi-fork simulations, wherein multiple candidate parameter sets are evaluated simultaneously. This multi-fork strategy expands parameter space exploration, yielding richer datasets that enable more accurate hypothesis refinement across iterations. Agents adaptively propose parameter updates based on intermediate simulation outcomes, enabling efficient navigation of complex trade-offs among structural, thermodynamic, and transport properties. The framework reproduces key experimental properties within 5% accuracy while maintaining consistency with atomistic reference behavior, achieving convergence without manual intervention. The modular architecture is readily extensible to other molecular systems and can accommodate additional targets, constraints, or simulation engines, providing a general agentic-AI platform for transferable CG model development.

**TOC GRAPHICS:** 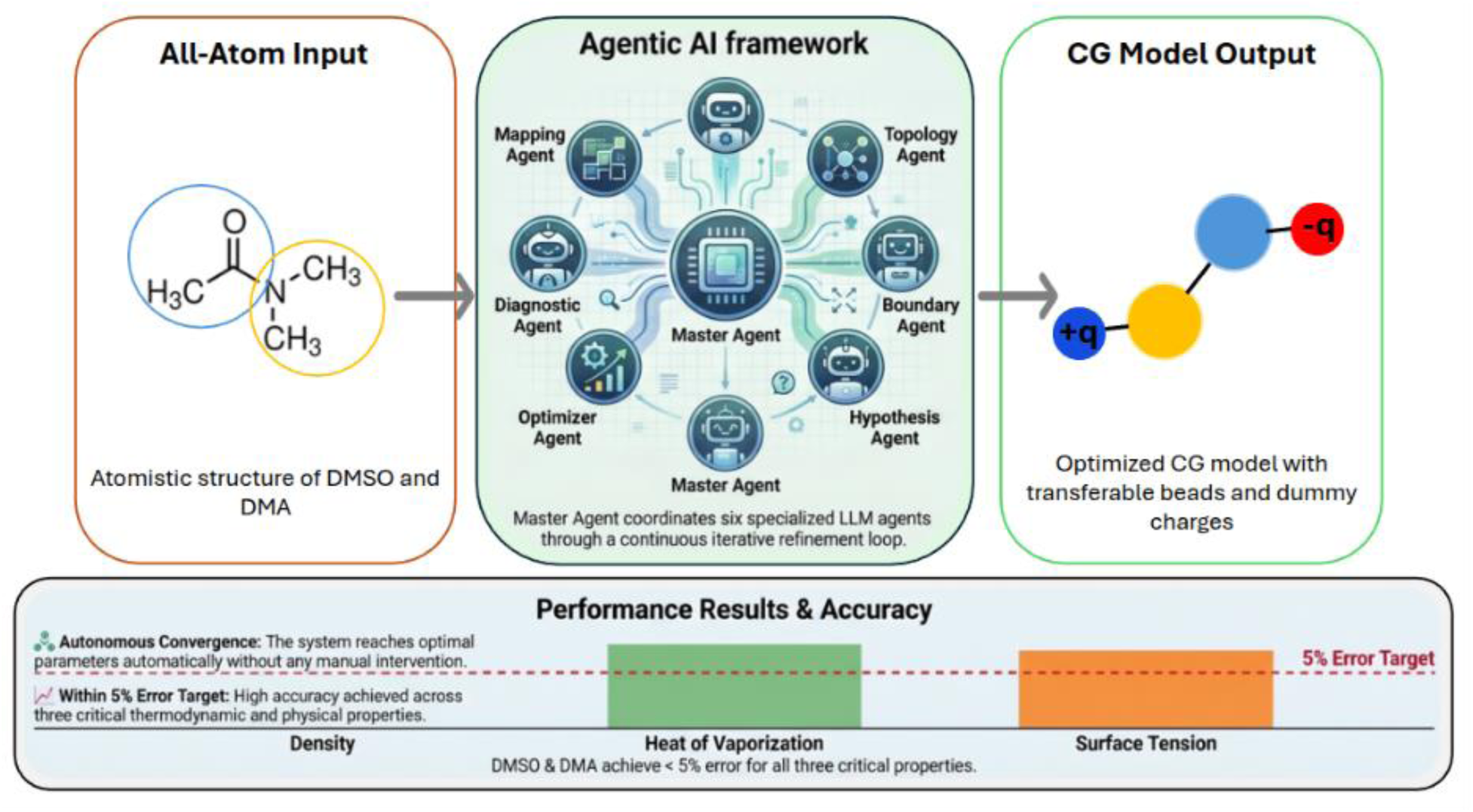

## INTRODUCTION

Coarse-grained (CG) molecular dynamics (MD) simulations have been used extensively for investigating molecular systems at mesoscopic scales with significantly lower computational cost compared to all-atom simulations^1–3^. By systematically reducing the degrees of freedom through mapping multiple heavy atoms onto single interaction sites (beads), CG models enable simulations that approach experimental length scales and access longer timescales while preserving essential physicochemical properties^4^. This computational efficiency is particularly valuable for studying processes such as self-assembly and the architectures of soft and hybrid materials, with applications ranging from pharmaceutical formulation to materials science^5,6^.

The development of CG models typically involves two key steps^3,7–15^. The first is the selection of a mapping scheme that defines how CG beads represent the underlying all-atom structure, and the second is the development of force-field (FF) parameters that describe bonded and non-bonded interactions between CG beads. Both steps present distinct challenges and are most often performed manually, relying heavily on the researcher’s experience and intuition. Mapping schemes are usually chosen based on atomic functionality or the need to preserve specific structural features. While this strategy is often effective, it can introduce downstream challenges during FF parameterization, as not all mapping choices permit the simultaneous reproduction of multiple target properties with acceptable accuracy.

To address these challenges, higher-dimensional optimization algorithms such as particle swarm optimization (PSO)^8,16,17^, Boltzmann inversion^18^, force matching^19^, and relative entropy minimization^20^ have been employed. In addition, machine learning (ML)-based approaches have increasingly been integrated with optimization algorithms to accelerate parameter exploration and improve model accuracy^9^. However, these ML-assisted methods typically inherit the same sequential structure as conventional workflows, treating mapping and FF parameterization as independent stages and therefore lacking the capacity to reason across their coupled dependencies. A deeper limitation is that the physically meaningful or high-performing regions of the joint mapping-parameter space are not known a priori, meaning that even sophisticated search strategies cannot efficiently identify which parameter ranges or local neighbourhoods’ merit exploration — particularly when the mapping scheme itself strongly governs what accuracy is achievable. Consequently, the conventional separation between mapping and parameter optimization limits the robustness and transferability of the resulting CG models and hinders the concurrent reproduction of multiple thermodynamic and structural properties.

These limitations motivate the need for a more integrated and autonomous approach to CG model development. Rather than treating mapping and FF parameterization as sequential and largely independent tasks, CG model construction is better viewed as a coupled and iterative decision-making problem, in which choices made at one stage strongly constrain what can be achieved at the next. Moreover, because promising regions of the combined mapping-parameter space are not known in advance, efficient model development requires repeated cycles of proposing, evaluating, rejecting, and refining candidate models. Thus, CG model development is fundamentally not only an optimization problem but also a hypothesis-driven search problem, which is particularly well suited for an automated agentic AI workflow, where specialized agents can coordinate reasoning and exploration across multiple interdependent tasks that are otherwise handled manually.

In recent years, the emergence of LLMs has created new opportunities for automating scientific workflows that traditionally required substantial domain expertise and human oversight^21,22^. Recent studies have demonstrated the ability of LLMs to assist with molecular design^23^, property prediction^24^, and scientific reasoning tasks^25^. Furthermore, agentic AI frameworks — in which multiple LLMs autonomously access external tools, execute specialized scripts, perform non-overlapping subtasks, and collaborate toward a shared goal — have demonstrated efficacy across diverse scientific and engineering domains, offering key advantages including enhanced error detection through mutual verification, improved task decomposition and parallel processing, and iterative output refinement through agent-to-agent feedback loops^26–28^. The application of agentic AI to molecular simulation and computational chemistry has begun to attract attention, though it remains nascent^29–32^. Early efforts demonstrated that LLM-based agents could autonomously execute quantum chemistry calculations and interpret their outputs, reducing the need for manual scripting and expert intervention^33^. More recently, multi-agent frameworks have been applied to automated FF development and simulation setup for small molecules and porous materials, where iterative feedback between simulation execution agents and parameter refinement agents has been shown to accelerate convergence toward target properties^21,34–36^. Agentic workflows have also been explored for protein structure prediction and de novo protein design, where agents combining knowledge retrieval, physics-based simulation, and results analysis have demonstrated the ability to autonomously design proteins with targeted mechanical properties^37,38^. At the materials scale, physics-aware multi-agent systems have been deployed for alloy design and discovery, autonomously performing atomistic simulations and iteratively refining hypotheses about composition–property relationships^29,35^.

To the best of our knowledge, however, the coupled problem of simultaneously optimizing CG mapping schemes and FF parameters through physically motivated hypothesis generation has not been addressed within an agentic AI framework, a critical gap given that mapping and parameterization decisions are deeply interdependent and that the combinatorial search space involved is too large and irregular for conventional optimization strategies alone. Additionally, LLMs are particularly suited to this role because, unlike traditional optimization methods such as Bayesian optimization^39,40^ or reinforcement learning^41^, they can incorporate domain knowledge, interpret intermediate simulation results in physical terms, and generate hypotheses that are both numerically constrained and chemically interpretable, a capacity for language-grounded physical reasoning that is essential when navigating the irregular, high-dimensional landscape of coupled mapping and FF parameter spaces.

In this work, we introduce an agentic AI framework, CGAgentX, in which specialized large language models (LLMs) function as autonomous agents that collaboratively explore mapping schemes and FF parameters within a closed-loop workflow. Specifically, using two widely used solvents, dimethyl sulfoxide (DMSO) and N,N-dimethylacetamide (DMA), as model systems, we demonstrate the framework’s ability to autonomously design viable CG mapping schemes while simultaneously optimizing the associated FF parameters. Both molecules are extensively characterized experimentally and computationally, and their broad relevance in biological and industrial applications makes them natural benchmarks for CG model validation. DMA and DMSO provide particularly stringent test cases because of (i) the presence of multiple plausible mapping schemes and (ii) their strong molecular polarity, which necessitates explicit treatment of electrostatic interactions through the placement of dummy beads.

## METHODS

CGAgentX comprises six specialized LLM agents — the Mapping Agent (MA), Topology Agent (TA), Boundary Agent (BA), Diagnostic Agent (DA), Hypothesis Agent (HA), and Optimizer Agent (OA) — each responsible for a distinct aspect of the CG model development workflow and equipped with a dedicated set of tools and scripts invoked through tool-calling capabilities. A Master Agent coordinates the sequential and iterative invocation of all agents, enforcing execution orders and managing data flow across the closed-loop workflow (see Supplementary Information for more details). All the agents were initialized using Moonshot AI’s Kimi K2.5 LLM model hosted on Virginia Tech ARC’s computational cluster.

The workflow begins with the MA, which is invoked only once at the start of the first epoch. If a CG mapping scheme already exists, either from a previous run or a user-supplied definition, the MA bypasses scheme generation, and the workflow proceeds directly with the existing mapping. Otherwise, it parses SMILES representations using an internal SMILESParser utility to extract atomic connectivity, bonding patterns, and connected components and uses these molecular features to validate and finalize a new CG grouping. Crucially, the MA also has access to a library of previously developed transferable CG beads, enabling it to incorporate established bead types into new mapping schemes rather than constructing every representation from scratch. The TA then orchestrates a multi-step system preparation pipeline, sequentially invoking a center-of-mass coarse-graining script to generate CG coordinates from the all-atom structure, Packmol^42,43^ to assemble a condensed-phase simulation box at the target experimental density, and PSFgen^44^ to produce NAMD-compatible topology files^45,46^. It additionally maps atomistic MD trajectories onto the CG representation via an in-house AA2CG conversion script, yielding a CG trajectory used for reference distribution analysis. The BA analyzes this trajectory to extract bond length, bond angle, and radial distribution function statistics, using these to establish physically informed initial parameter bounds; in the absence of atomistic reference data, it estimates bead sizes from radius-of-gyration calculations using internal geometry routines.

With the CG system prepared and parameter bounds established, the framework enters its iterative optimization loop. At each iteration, the HA formulates a single physically motivated hypothesis for the next parameter adjustment, drawing on the diagnostic report and performance scores from all parallel forks. The OA then translates this hypothesis into nfork (2, 4, or 8) distinct parameter sets, each representing a different concrete realization of the same underlying hypothesis, and distributes them across parallel simulation forks so that each fork explores a different region of the parameter space consistent with the same physical rationale. Once all forks have completed, the DA collects and analyzes the full set of results, evaluating system phase behavior, distinguishing liquid, solid, and collective flow regimes, simulation stability, and thermodynamic property deviations using standard Linux text-processing utilities and Python codes, and contextualizing its assessment against optimization history loaded from a structured memory directory. The DA produces a diagnostic report that includes performance scores, warnings, and recommendations for parameter boundary revision communicated to HA and BA. The HA then reads the scores from all forks together with the diagnostic report to formulate an updated hypothesis, after which the OA generates revised parameter sets for the next round of forks. Both agents read and write structured JSON memory files and log into their full prompts for reproducibility and debugging. This closed-loop architecture ensures that hypothesis generation is always informed by the collective behavior of all parallel parameter evaluations rather than any single simulation outcome.

To systematically explore the space of possible CG representations, we performed three independent runs of the full workflow for both DMSO and DMA. After the first run generated a mapping scheme, that scheme was added to the existing library, and the MA was instructed to exclude it from future generations; the same procedure was repeated after the second run so that by the third run, two schemes were already cataloged. This sequential exclusion ensured that the MA was forced to propose a genuinely new CG representation in each successive run. For each mapping scheme, three independent optimization runs were further performed per fork setting to evaluate the effect of different fork counts on optimization efficiency.

To build CG models with intrinsic temperature transferability, the optimization was performed simultaneously at two temperatures for each solvent — 298 K and 323 K for DMSO, and 298 K and 313 K for DMA — so that the fitness function evaluated property deviations across both thermodynamic states at every iteration, forcing the agents to identify parameter sets that capture physically meaningful temperature dependence rather than overfitting to a single state point. Once the optimization runs converged, the best-performing parameter set from each fork and mapping scheme combination was carried forward into an independent 100 ns production simulation to validate the optimized models against target structural and thermodynamic properties under extended sampling conditions. During the validation simulation, the thermodynamic properties were derived from the final 30 ns of trajectory data. Four thermodynamic target properties, density, heat of vaporization, surface tension, and dipole moment, were evaluated at two temperatures per solvent: 298 K and 313 K for DMA (experimental targets: density = 0.9361 and 0.9241 g/cm³, Hvap = 10.951 and 10.743 kcal/mol, ST = 32.43 and 31.56 dyn/cm, dipole = 3.72 D)^47–49^ and 298 K and 323 K for DMSO (experimental targets: density = 1.0950 and 1.0702 g/cm³, Hvap = 12.648 and 12.400 kcal/mol, ST = 42.09 and 40.05 dyn/cm, dipole = 3.96 D)^50–52^. Three independent optimization runs were conducted per solvent, each exploring three fork counts (nfork = 2, 4, and 8) across all three mapping schemes, yielding a total of 54 independent optimizations (3 runs × 3 forks × 3 schemes × 2 solvents). The details of the simulation setup are included in the supplementary information. All optimizations were performed on AMD EPYC 7702 CPU cores in a parallelized framework, with each forked simulation utilizing 16 CPU cores. Validation simulations of 100 ns were executed on NVIDIA A30 GPUs.

## RESULTS AND DISCUSSION

Applying the agentic framework to DMSO and DMA, the MA autonomously generated three distinct CG representations for each solvent across the three independent workflow runs, shown in **Fig. 2**, drawing on both its molecular parsing capabilities and the library of transferable CG beads. For DMSO, the proposed mappings ranged from a minimal coarse-graining, in which a central sulfur atom and one methyl group were grouped into a single MS2 bead while the oxygen atom and the second methyl group formed the OCB bead in Scheme 1, to more detailed representations incorporating two charged dummy beads to explicitly capture electrostatic interactions (Schemes 2 and 3). For DMA, the agent suggested a more atomistic representation in Scheme 3, which included three CG beads and two charged dummy beads. By leveraging the transferable bead library, the MA reused the MS2 bead from the amino acid model^10^ in Schemes 1 and 2 for DMSO and the CGD2 bead from the DMF solvent model in Scheme 3 for both DMSO and DMA, ensuring compatibility with existing CG models developed in our group. Across all proposed mappings, the agent consistently preserved the polar character and dipole direction of both molecules through explicit dummy bead placement, indicating that the agent’s reasoning prioritized electrostatic fidelity alongside structural simplification. Moreover, the connectivity-aware validation by MA ensured that all proposed mappings maintained molecular integrity by preventing physically invalid groupings of disconnected atoms.

**Figure 1:**
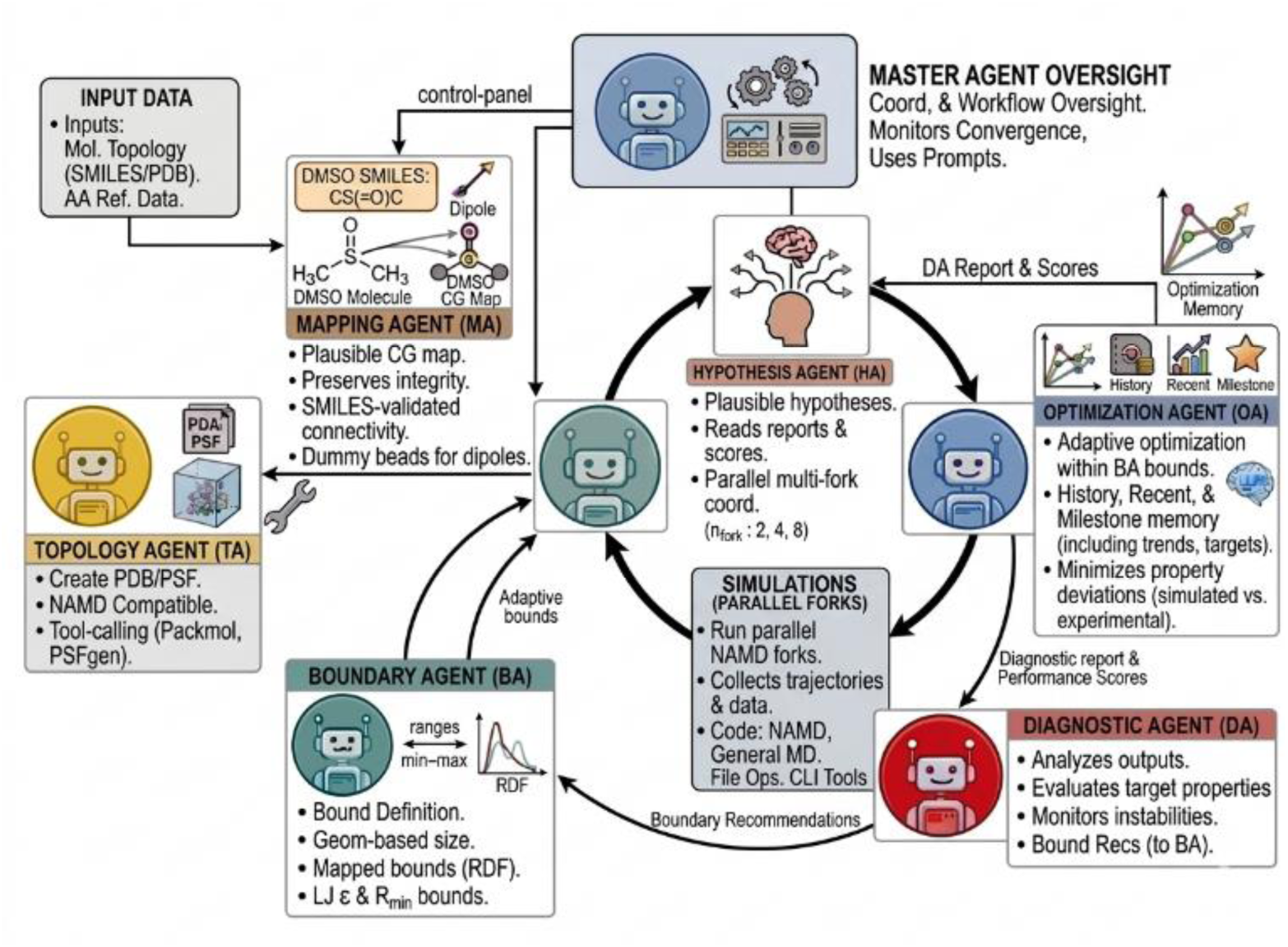
The CGAgentX framework with six specialized agents with tool-calling capabilities is used for autonomously developing DMSO and DMA coarse-grained models. The MA processes the all-atom SMILE/PDB input along with the experimental targeted properties to generate CG mapping schemes. The TA and BA incorporate the generated mapping schemes to prepare the CG systems for MD simulations. The HA proposes plausible hypotheses for OA to generate parameters to test using a multi-fork setup. All agents utilize centralized memory and learn from previous successes and failures, achieving the best parameter that yields property values close to the experimental range.

**Figure 2:**
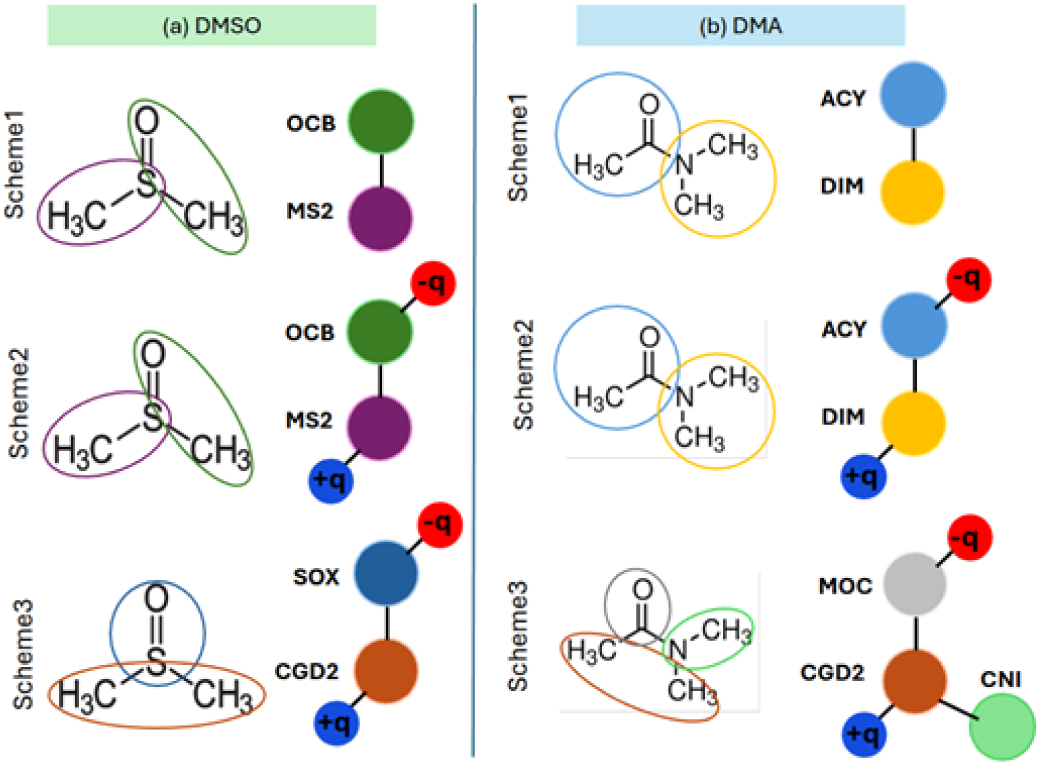
Three different CG mapping schemes proposed by the mapping agent for (a) DMSO and (b) DMA. The blue and red beads indicate the positively and negatively charged dummy beads, respectively.

With each mapping scheme established, the framework entered its iterative optimization loop, in which the HA, OA, and DA operated in concert to refine FF parameters. The HA generated physically motivated hypotheses at each iteration, using LLM-based chemical reasoning informed by previous epochs’ results, diagnostic reports, and system-specific chemical details. A central question for any such agent is whether it engages with the underlying physics of the system or merely searches for parameter combinations that reduce a numerical score. To address this question, we systematically analyzed the scientific rationale field of every hypothesis produced by the HA for all schemes’ optimization and 54 full runs, covering three replicates at each of three fork counts, yielding 3073 hypotheses (1528 in DMA and 1545 in DMSO) in total.

The rationales are not simple action-value statements: 73% contain at least one explicit mechanism-level reasoning signature shown in **Supplementary Figure S2(a)** for Scheme 3. The most prevalent signatures are compensation moves, in which the agent balances a perturbation in one parameter against a physically motivated counter-perturbation in another (38% of the hypotheses), and direct invocation of the physical relationship, such as μ = q × d, that couples the molecular dipole moment to the dummy-bead charge magnitude and separation distance (37%). A further 16% of hypotheses invoke by name the geometric constraint that charged dummy beads must reside within their parent bead’s Lennard-Jones σ/2 radius; 13% contain explicit mechanistic diagnoses of simulation crashes (e.g., identifying electrostatic catastrophe arising from dummy beads that carry charges in the absence of van der Waals repulsion); and 10% recognize that a parameter has reached its physical or numerical boundary and explicitly pivot to an alternative physical lever to achieve the same optimization target. For schemes 1 and 2, a similar trend is observed as for scheme 3 (see Supplementary Information for scheme-specific breakdowns). These signatures are non-exclusive and frequently co-occur within a single rationale, consistent with the multi-objective character of the FF refinement problem.

The character of this reasoning is best illustrated by three representative cases reproduced in **Supplementary Figure S2(b)**. In case (b1), corresponding to DMSO Scheme 3, Run 1, Fork 2, iteration 89, the HA simultaneously reduces dummy-bead charges to lower the electrostatic contribution to surface tension, which scales as q², and extends the dummy-bead bond lengths to preserve the molecular dipole moment through the identity μ = q × d, while also increasing the Lennard-Jones ε parameter to recover the cohesive energy lost from the bulk due to the charge reduction. In case (b2), DMA Scheme 3, Run 3, Fork 2, iteration 28, the agent identifies that LJ ε has been driven to its upper physical boundary and, rather than continuing to apply incremental pressure against this constraint, redirects its optimization effort to the electrostatic and bond-geometry subsystems to address the remaining heat-of-vaporization and surface-tension errors through an independent physical mechanism. In case (b3), DMSO Scheme 3, Run 1, Fork 2, iteration 6, the agent responds to a velocity-overflow crash by first diagnosing the underlying mechanism and then constructing a coupled corrective hypothesis that reduces dummy-bead charges, stiffens core bonds to suppress overlap, and compensates for the charge reduction through geometric adjustments that preserve the target dipole moment. The detailed breakdown of behavioral metrics and per-run breakdowns are shown in **Supplementary Figures S3-S6**. In each case, the hypothesis engages a named physical relationship or constraint, identifies trade-offs between coupled target properties, and constructs a multi-parameter move that respects them, a proof of mechanism-level reasoning rather than an undirected parameter search.

To test each hypothesis, the OA translated it into nfork distinct parameter sets, and all simulations were conducted in parallel within a closed feedback loop. **Fig. 3** demonstrates this process for DMSO Scheme 2, Run 1, with four forks. In the first 10 epochs, the error steadily decreased to less than 8%, as is evident in **Fig. 3 (i)-(j)**. In epoch 15, a transient spike in the composite percentage error was observed (from 87.79 to 152.73), triggered by an aggressive corrective measure taken by the HA agent, which increased the LJ epsilon and dummy bead charge in response to the underestimation of all the thermodynamical properties encountered in the previous epoch. In the next epoch, 16, the HA correctly identified over-cohesion (overestimating Hvap by 45% and surface tension by 85–96%) and lowered the epsilon and charge magnitude, which in turn lowered the composite error. This adaptive reasoning capability of the HA agent mirrored the corrective intuition of an experienced researcher, rather than propagating a failed parameter set. After 50 epochs, this process yielded the best parameter set for DMSO Scheme 2, achieving a 1.71% error. During optimization, the parameter bounds progressively contracted toward physically relevant regions shown in **Supplementary Figure S13**, reflecting the exclusion of unproductive regions of parameter space and convergence toward optimal solutions. While convergence trends are consistent across independent runs, minor variations in early-stage trajectories reflect inherent stochastic differences in parameter initialization by LLM agents, highlighting the importance of multi-run averaging for robust assessment of optimization performance.

**Figure 3:**
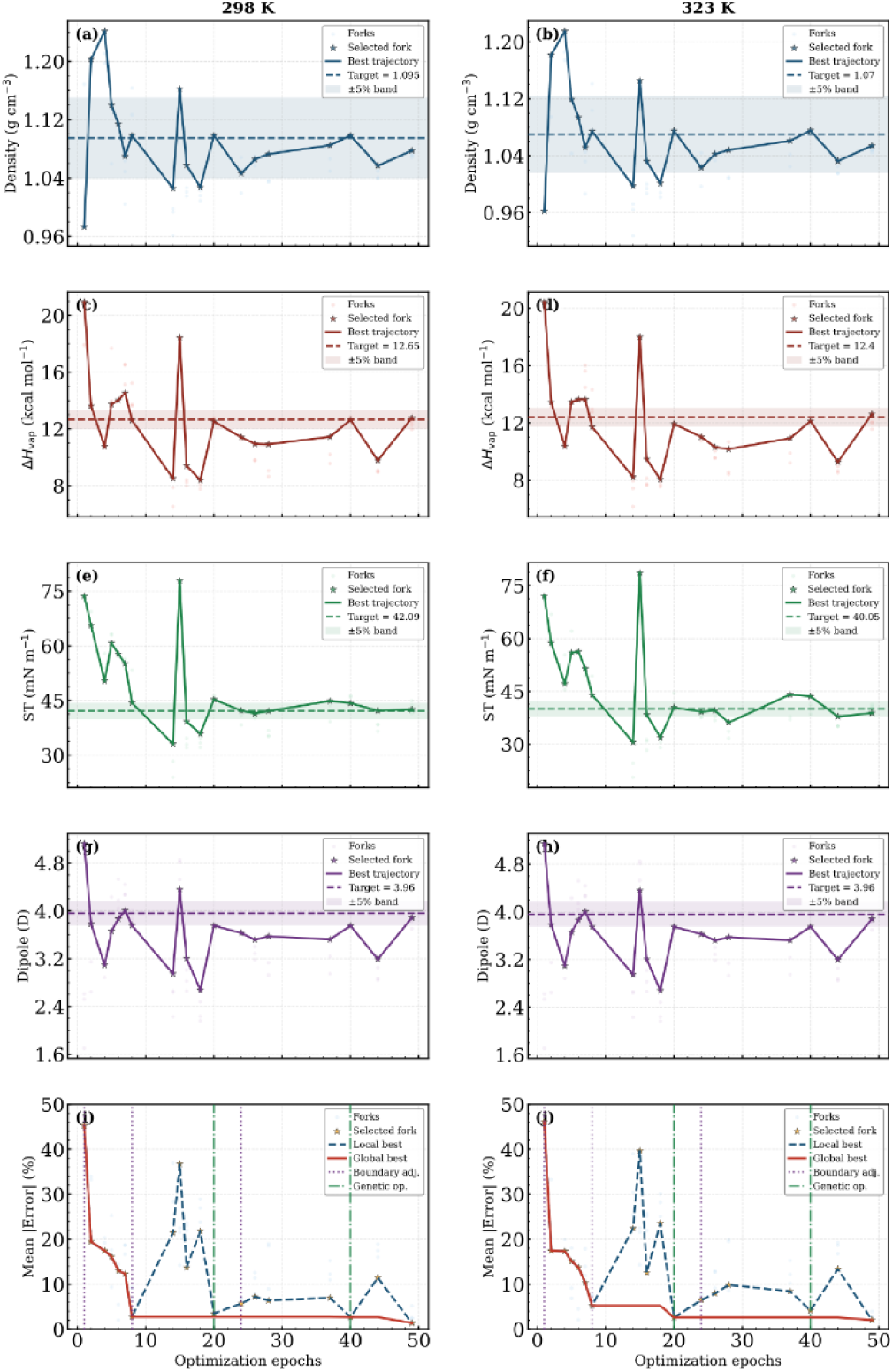
Multi-fork optimization with nfork=4 for mapping scheme 2 of DMSO at 298K and 323K, respectively. (a)-(d) show the evolution of density, heat-of-vaporization, surface tension, and dipole moment with the optimization epochs. The dashed lines indicate the experimental targets, along with a 5% range in shades. (i)-(j) show the corresponding percentage error per property of the best fork in each epoch by the blue dashed line, and the global best is indicated by the solid red line.

The properties and their percentage errors obtained during parameter development for two different temperatures with three different mapping schemes and forks are shown in **Supplementary Table S2**. Across three independent optimization runs, the best parameters for DMSO were obtained at Run 3, Fork 8, and Scheme 3, yielding a 1.6% mean error across properties, whereas for DMA Run 3, Fork 8, and Scheme 1, 0.2% error was achieved (see **Supplementary Table S3**).

### Effect of number of forks

Regardless of the mapping scheme, more forks accelerated convergence, as shown in **Fig. 4** for Run 1 and in **Supplementary Figures S14-S16** for all runs. Overall, Fork 8 reached <10% mean error by epoch 8 versus epochs 12 and 21 on average for Forks 4 and 2, respectively. The hypotheses are tested more rigorously *via* simultaneous simulations with an increasing number of forks, enabling border explorations in a few epochs, which led to the initial larger drop while having multiple smaller drops in later epochs evident from Fork 8. This acceleration reflects a qualitative enrichment in HA-generated hypotheses as fork count increases. Analysis of the HA’s reasoning text reveals that the rate at which hypotheses explicitly ground proposals in the DA’s aggregated report rises from 65.8% to 90.4% (DMSO) and 79.6% to 93.8% (DMA) going from Fork 2 to Fork 8 (see Supplementary Information for a more detailed breakdown). Simultaneously, the frequency of hypotheses citing specific numerical residual errors increases from 48.3% to 82.6% in DMSO, indicating a shift from qualitative direction-only adjustments toward quantitatively anchored reasoning. The average number of distinct “sub-strategy markers” per hypothesis — which approximate the number of distinct exploration directions that the HA is simultaneously proposing to OA — scales with fork count (3.01 at Fork 2 versus 4.36 at Fork 8), reflecting the HA’s deliberate decomposition of each proposal into orthogonal exploration directions matched to the available fork budget. Collectively, a larger fork ensemble furnishes the DA with a more statistically representative picture of the parameter landscape, enabling richer diagnoses that the HA converts into more guided, numerically grounded hypotheses with a compounding feedback effect that leads to faster convergence for a greater number of forks.

**Figure 4:**
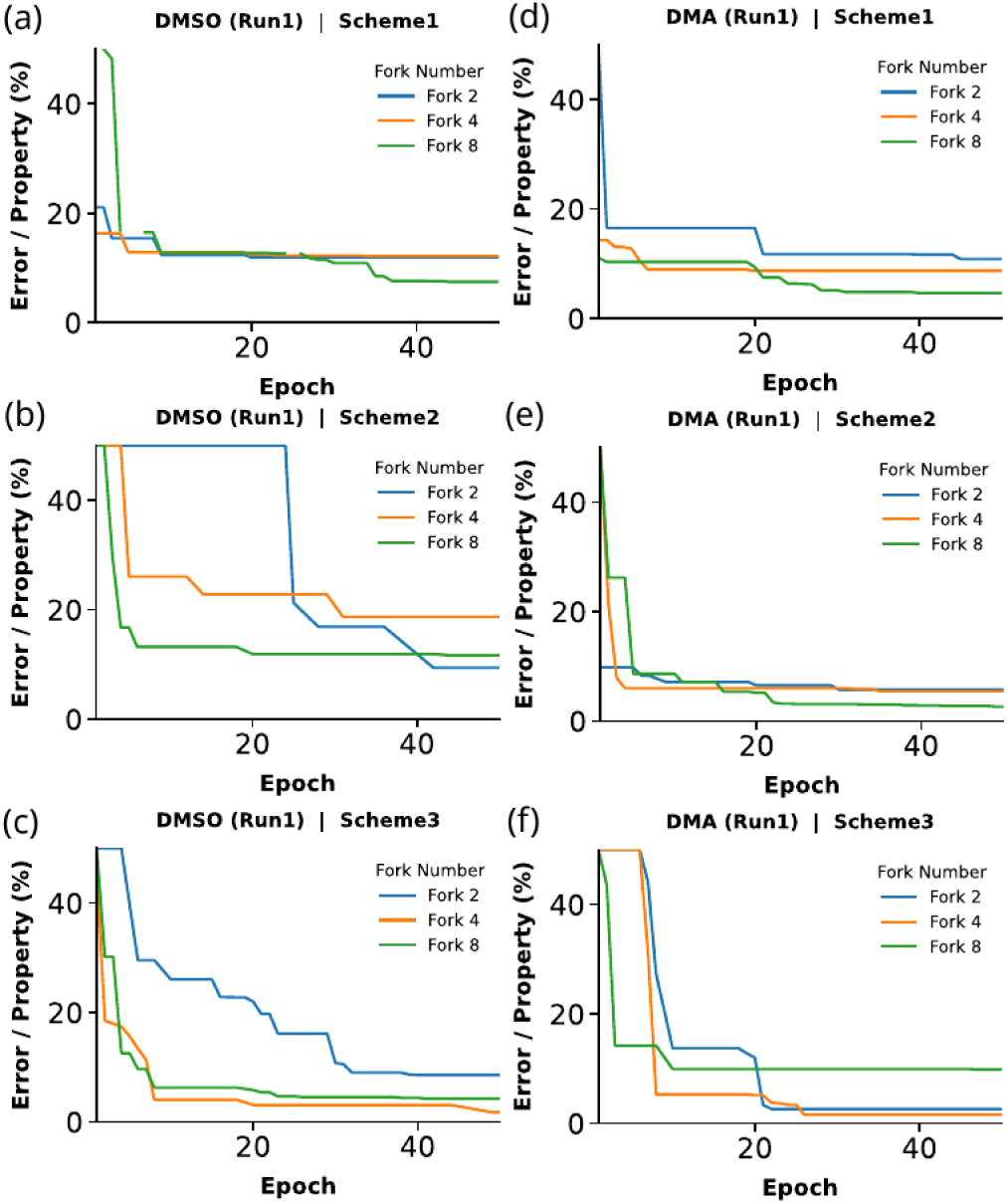
The percentage error per property of the global best as a function of optimization epochs at 298 K for (a)-(c) DMSO and (d)-(f) DMA for three different mapping schemes. Forks 2, 4, and 8 are denoted by blue, orange, and green lines, respectively.

### Effect of the mapping scheme

Schemes 2 and 3, which incorporate charged dummy beads to capture electrostatic interactions, yielded mean errors of 2.6% for DMA and 4.2% for DMSO across the best optimization runs as reported in **Supplementary Table S4**. In contrast, Scheme 1, which lacks explicit electrostatic representation, produced mean errors above 6.5%, driven primarily by the underestimation of heat of vaporization (see **Supplementary Table S5, Supplementary Figure S17**). Heat of vaporization constitutes the dominant source of error in Scheme 1 for both DMSO and DMA, contributing error margins of 15% and 8%, respectively. The absence of dummy charged beads in Scheme 1 results in weaker cohesive interactions among solvent molecules, which leads to an underestimation of the heat of vaporization in both DMSO and DMA, as well as the density in DMSO. The introduction of dummy beads in Schemes 2 and 3 improved the heat of vaporization accuracy, reflecting the importance of explicit electrostatic representation in capturing intermolecular cohesion at the CG level.

### Transferability across temperatures

Across all schemes, the mean percentage errors correlate across temperatures, with the best parameter sets showing temperature transferability and only modest changes in mean error across the 25 K and 15 K temperature intervals for DMSO and DMA, respectively, corresponding to an average temperature sensitivity of approximately 0.05% per Kelvin. For both temperatures, all the properties were observed within 7% of the experimental target. No systematic degradation in accuracy is observed at higher temperatures, indicating that the optimized parameters capture physically meaningful temperature dependence rather than overfitting to a single thermodynamic state. Textual analysis of the HA’s reasoning further corroborates this transferability. In Scheme 3 DMSO runs, explicit numerical temperature references in the hypotheses increase monotonically with the fork count, rising from 6% at Fork 2 to 26% at Fork 8. Similarly, the rate at which the agent explicitly flags single-temperature residual mismatches increases from 4% to 21% over the same range, reflecting the richer per-iteration feedback available at higher fork counts (see **Supplementary Figures S7-S12**). This effect is present, albeit weakly, across the other mapping schemes and both solvents. For “temperature-gap events” — where a target property is within tolerance at one temperature but not the other — the DMSO Fork 8 HA explicitly addresses the off-target state in 41.7% of cases, compared to only 6.7% at Fork 2. The resulting best-fork parameter sets achieve comparable accuracy at both state points for every target property, with mean temperature sensitivities below 0.1%/K for bulk properties. These findings confirm that the two-temperature optimization constraint propagates effectively through the HA’s reasoning, yielding temperature-transferable parameter sets across all mapping schemes and both solvents.

### Solvent-specific effects

DMSO is more challenging than DMA for CG modeling due to its stronger electrostatic character (dipole moment = 3.96 D) and higher surface tension (ST > 40 dyn/cm). The surface tension is overestimated in Fork 2 and Fork 4 for schemes 1 and 3. This persistent overestimation reflects a fundamental challenge: surface tension is a collective interfacial property arising from the balance of cohesive forces at the liquid–vapor interface and is sensitive to lack of hydrogen-bond-like^53,54^ interactions, which suppresses interfacial density fluctuations and capillary wave contributions^55,56^. Even all-atom FFs exhibit this behavior; CHARMM36 yields a surface tension of 44.36 dyn/cm for DMSO, 6% above experiment (see **Supplementary Table S1**), indicating that this property is fundamentally difficult to capture.

Three 100 ns validation simulations conducted with the best parameters obtained from the optimization cycles yield a closer match to experimental reference values, demonstrating the reliability and transferability of the CGAgentX framework. For the best DMA result (Scheme 1, Fork 8), the mean errors were 0.77 ± 0.39% and 1.17 ± 0.74% at 298 K and 313 K, respectively, while for the best DMSO result (Scheme 3, Fork 4), the mean errors were 1.61 ± 0.02% and 2.54 ± 1.04% at 298 K and 323 K (**Table 1**, **Supplementary Tables S2, S6–S7**), within a 1% range of the optimization cycle errors. The consistency of these errors across temperatures confirms that the optimized parameters are transferable beyond optimization cycles and that their accuracy is preserved over significantly longer simulation timescales.

**Table 1:**
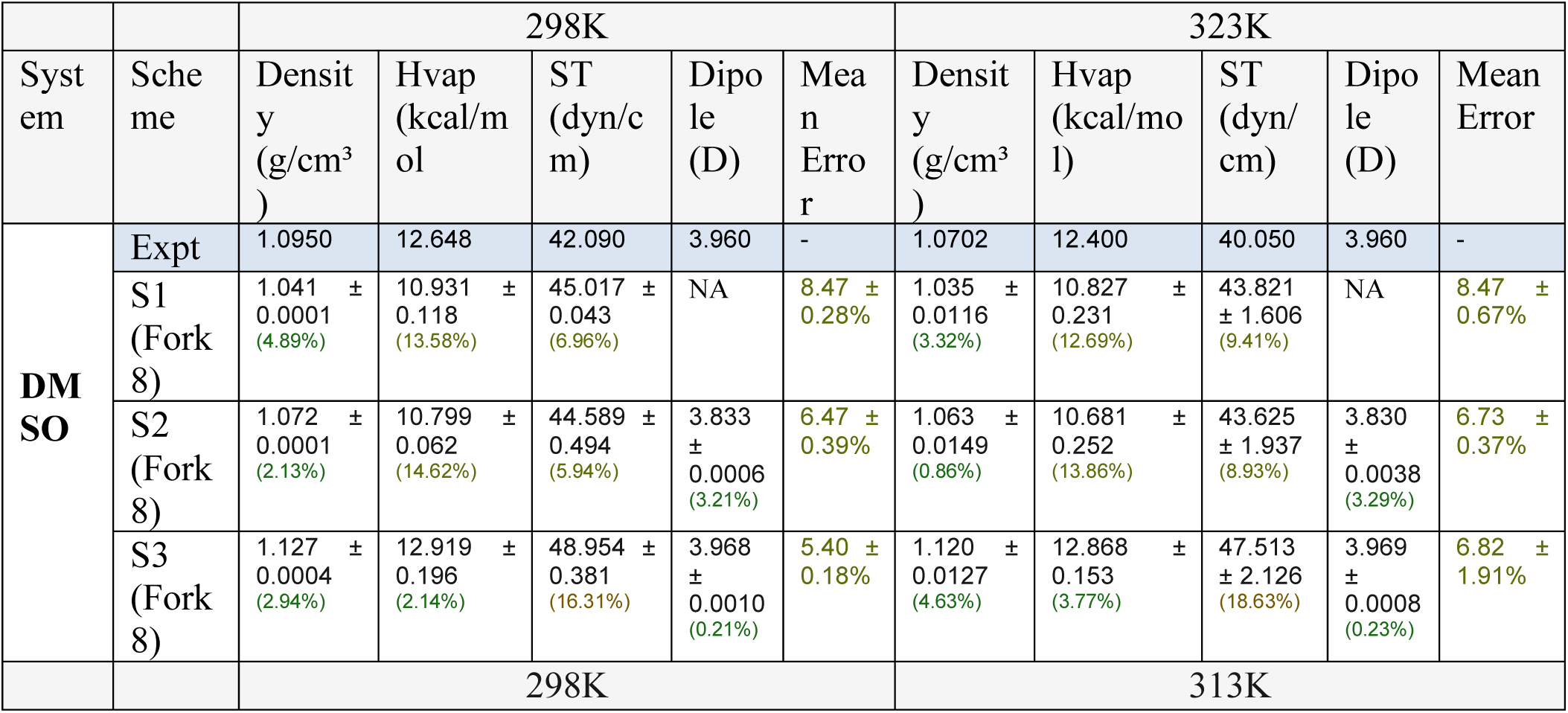
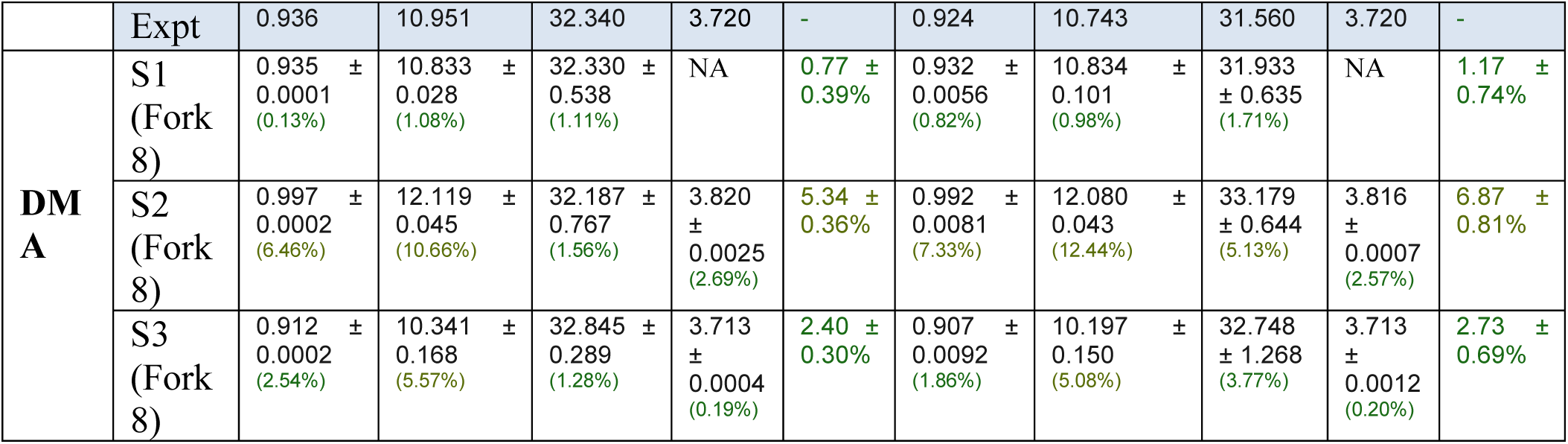
DMSO and DMA properties after 100 ns of MD simulation starting from the best optimized parameters by eight forks for three different mapping schemes at two different temperatures. The property values are averaged over three independent runs, and the percentage error for each of the properties is shown within brackets. The dipole moment cannot be obtained from Scheme 1 due to the absence of charged dummy beads and is denoted as not applicable (NA).

|  |  | 298K |  |  |  |  | 323K |  |  |  |  |
| --- | --- | --- | --- | --- | --- | --- | --- | --- | --- | --- | --- |
| Syst<br>em | Sche<br>me | Densit<br>y<br>(g/cm<sup>3</sup><br>) | Hvap<br>(kcal/m<br>ol) | ST<br>(dyn/c<br>m) | Dipo<br>le<br>(D) | Mea<br>n<br>Erro<br>r | Densit<br>y<br>(g/cm<sup>3</sup><br>) | Hvap<br>(kcal/mo<br>l) | ST<br>(dyn/<br>cm) | Dipo<br>le<br>(D) | Mean<br>Error |
| **DM**<br>**SO** | Expt | 1.0950 | 12.648 | 42.090 | 3.960 | - | 1.0702 | 12.400 | 40.050 | 3.960 | - |
|  | S1<br>(Fork<br>8) | 1.041 ±<br>0.0001<br>(4.89%) | 10.931 ±<br>0.118<br>(13.58%) | 45.017 ±<br>0.043<br>(6.96%) | NA | 8.47 ±<br>0.28% | 1.035 ±<br>0.0116<br>(3.32%) | 10.827 ±<br>0.231<br>(12.69%) | 43.821<br>± 1.606<br>(9.41%) | NA | 8.47 ±<br>0.67% |
|  | S2<br>(Fork<br>8) | 1.072 ±<br>0.0001<br>(2.13%) | 10.799 ±<br>0.062<br>(14.62%) | 44.589 ±<br>0.494<br>(5.94%) | 3.833<br>±<br>0.0006<br>(3.21%) | 6.47 ±<br>0.39% | 1.063 ±<br>0.0149<br>(0.86%) | 10.681 ±<br>0.252<br>(13.86%) | 43.625<br>± 1.937<br>(8.93%) | 3.830<br>±<br>0.0038<br>(3.29%) | 6.73 ±<br>0.37% |
|  | S3<br>(Fork<br>8) | 1.127 ±<br>0.0004<br>(2.94%) | 12.919 ±<br>0.196<br>(2.14%) | 48.954 ±<br>0.381<br>(16.31%) | 3.968<br>±<br>0.0010<br>(0.21%) | 5.40 ±<br>0.18% | 1.120 ±<br>0.0127<br>(4.63%) | 12.868 ±<br>0.153<br>(3.77%) | 47.513<br>± 2.126<br>(18.63%) | 3.969<br>±<br>0.0008<br>(0.23%) | 6.82 ±<br>1.91% |
|  |  | 298K |  |  |  |  | 313K |  |  |  |  |

|  | Expt | 0.936 | 10.951 | 32.340 | 3.720 | - | 0.924 | 10.743 | 31.560 | 3.720 | - |
| --- | --- | --- | --- | --- | --- | --- | --- | --- | --- | --- | --- |
| **DM**<br>**A** | S1<br>(Fork<br>8) | 0.935 ±<br>0.0001<br>(0.13%) | 10.833 ±<br>0.028<br>(1.08%) | 32.330 ±<br>0.538<br>(1.11%) | NA | 0.77 ±<br>0.39% | 0.932 ±<br>0.0056<br>(0.82%) | 10.834 ±<br>0.101<br>(0.98%) | 31.933<br>± 0.635<br>(1.71%) | NA | 1.17 ±<br>0.74% |
|  | S2<br>(Fork<br>8) | 0.997 ±<br>0.0002<br>(6.46%) | 12.119 ±<br>0.045<br>(10.66%) | 32.187 ±<br>0.767<br>(1.56%) | 3.820<br>±<br>0.0025<br>(2.69%) | 5.34 ±<br>0.36% | 0.992 ±<br>0.0081<br>(7.33%) | 12.080 ±<br>0.043<br>(12.44%) | 33.179<br>± 0.644<br>(5.13%) | 3.816<br>±<br>0.0007<br>(2.57%) | 6.87 ±<br>0.81% |
|  | S3<br>(Fork<br>8) | 0.912 ±<br>0.0002<br>(2.54%) | 10.341 ±<br>0.168<br>(5.57%) | 32.845 ±<br>0.289<br>(1.28%) | 3.713<br>±<br>0.0004<br>(0.19%) | 2.40 ±<br>0.30% | 0.907 ±<br>0.0092<br>(1.86%) | 10.197 ±<br>0.150<br>(5.08%) | 32.748<br>± 1.268<br>(3.77%) | 3.713<br>±<br>0.0012<br>(0.20%) | 2.73 ±<br>0.69% |

The density prediction of the best DMSO model (Fork 4 Scheme 3) can be benchmarked against both the experimental values and the SDK-based two-bead CG model of Shobhna et al^54^. At 298 K, the SDK-based model overestimates density by ∼0.64% (1.102 vs. 1.095 g/cm³), while our model underestimates it by ∼1.58% (1.0774 g/cm³). At 323 K, deviations remain similarly modest: ∼0.36% high for Shobhna et al. (1.074 g/cm³) versus ∼1.5% low for our model (1.0541 g/cm³), against the experimental target 1.0702 g/cm³. Critically, while Shobhna et al. optimized exclusively against density and membrane/PMF observables, the agentic AI-driven models were simultaneously parameterized to reproduce density, Hvap, surface tension, and dipole moment. The slightly larger density deviation is therefore an expected consequence of this more rigorous multi-property optimization, which confers broader thermodynamic transferability.

Structural validation against all-atom mapped reference trajectories further confirms the physical soundness of the optimized models (**Supplementary Figs. S18–S19**). For DMSO scheme 3, both the CG bond distributions and radial distribution functions (RDFs) of the core beads show close agreement with the all-atom mapped trajectories at both 298 K and 323 K. The RDF peaks and overall structure are well preserved within 12 Å, indicating that the framework successfully captures the local packing structure of DMSO in the CG representation. For DMA Scheme 3 (using the three-bead CGD2-CNI-MOC representation), the bond distributions are well reproduced, but the CGD2–CNI–MOC angle distribution is approximately 30% larger in the CG model compared to the all-atom mapped reference. This angular discrepancy arises because the bonded interactions in the CG model were parameterized jointly with nonbonded terms, which may sacrifice exact structural reproduction to optimize the aggregate thermodynamic fitness. The fitness function, which weighs thermodynamic properties equally, does not explicitly penalize structural deviations (see Supplementary Information). This represents a known trade-off in CG parameterization: bottom-up approaches that target structural reproduction (force-matching, iterative Boltzmann inversion) often yield poor thermodynamics, while top-down approaches that target thermodynamic properties may produce structural deviations^57^.

These results demonstrate that within a given mapping, the HA performs sophisticated mechanism-level physical reasoning by invoking parameter-named relationships, recognizing geometric constraints, and constructing multi-parameter compensation moves to efficiently traverse complex parameter space. Critically, increasing parallel forks enriches this reasoning through broader feedback, accelerating convergence by up to 2.6-fold; mapping sets the destination, hypothesis quality charts the path, and fork count drives the speed. Together, these capabilities position CGAgentX as a powerful and general framework for automated CG model development across diverse materials and biomolecules.

## CONCLUSION

The results reported here demonstrate that an autonomous multi-agent framework can successfully develop coarse-grained (CG) models for polar solvents without manual intervention, achieving an agreement with experimental targets within ∼5% across density, heat of vaporization, surface tension, and dipole moment while maintaining consistency with atomistic reference behavior across temperatures.

Taken together, the DMSO and DMA case studies identify that within a given mapping scheme, the Hypothesis Agent navigates the parameter space through mechanism-level physical reasoning rather than undirected searching. And increasing the number of parallel simulation forks amplifies this reasoning by providing richer feedback per hypothesis, accelerating convergence by up to 2.6-fold.

Beyond the specific systems studied, CGAgentX’s modular architecture is readily extensible to other solvents and complex molecular systems and can accommodate additional target properties, constraints, or simulation engines. By combining autonomous execution with transparent, hypothesis-driven decision-making, the framework provides interpretable rationales for parameter choices that can inform future FF development strategies — establishing a general agentic AI platform for transferable CG model development across a wide range of molecular systems.

Ultimately, the success of CGAgentX’s framework on these test systems points toward a generalizable paradigm for autonomous CG model development, with implications for accelerating molecular simulation workflows across a broad range of chemically and structurally diverse systems.

## SUPPORTING INFORMATION

All the codes used in the study are available to download from GitHub [[Link](https://github.com/Deshmukh-Group/Agentic-AI_CG_Solvent_Model)] and a dashboard to visualize the data [[Link](https://deshmukh-group.github.io/Agentic-AI_CG_Solvent_Model/)].

## AUTHOR CONTRIBUTIONS

SD conceptualized the idea. SS translated it into code, set up the framework, and ran all the simulations. Both authors contributed to manuscript writing and data analysis.

## ACKNOWLEDGEMENTS

The authors thank the Advanced Research Computing Center at Virginia Tech for providing computational resources and technical support that have contributed to the results reported within this study. This work was supported by GlycoMIP, a National Science Foundation (NSF) Materials Innovation Platform (MIP) funded through Cooperative Agreement DMR-1933525. Authors acknowledge the usage of AI platforms (Claude and Gemini models) for refining the TOC and schematic figures.

## NOTES

There are no conflicts to declare.

