## Supplementary Information for "CGAgentX: Agentic AI Framework to Develop Transferable Coarse-Grained Models"

### Agentic-AI Framework for All-Atom Simulation Setup

In traditional MD setups, researchers manually perform tasks such as molecular packing, topology generation, and simulation configuration in sequential order, which can be error-prone and time-intensive, especially for all-atom systems involving large biomolecules or solvents. The agentic AI framework for an all-atom simulation setup presents a transformative approach to minimizing and rectifying errors by leveraging large language models (LLMs) in loops as autonomous decision-makers. Agentic AI solves these challenges by invoking modular agents that interact dynamically, diagnose issues, and execute corrections with minimal human intervention. Our framework demonstrates all-atom equilibrium MD simulations set up for polar solvents DMSO and DMA using the NAMD[1] engine with the CHARMM36 force field[2]. By integrating five specialized agents for planning (Planner Agent), packing solvent molecules into a cubic simulation box (Packing Agent), topology assignment (Topology Agent), simulation preparation (Simulation Agent), and diagnostics (Diagnostic Agent), the workflow streamlines the generation of production-ready simulation environments shown in **Figure S1**.

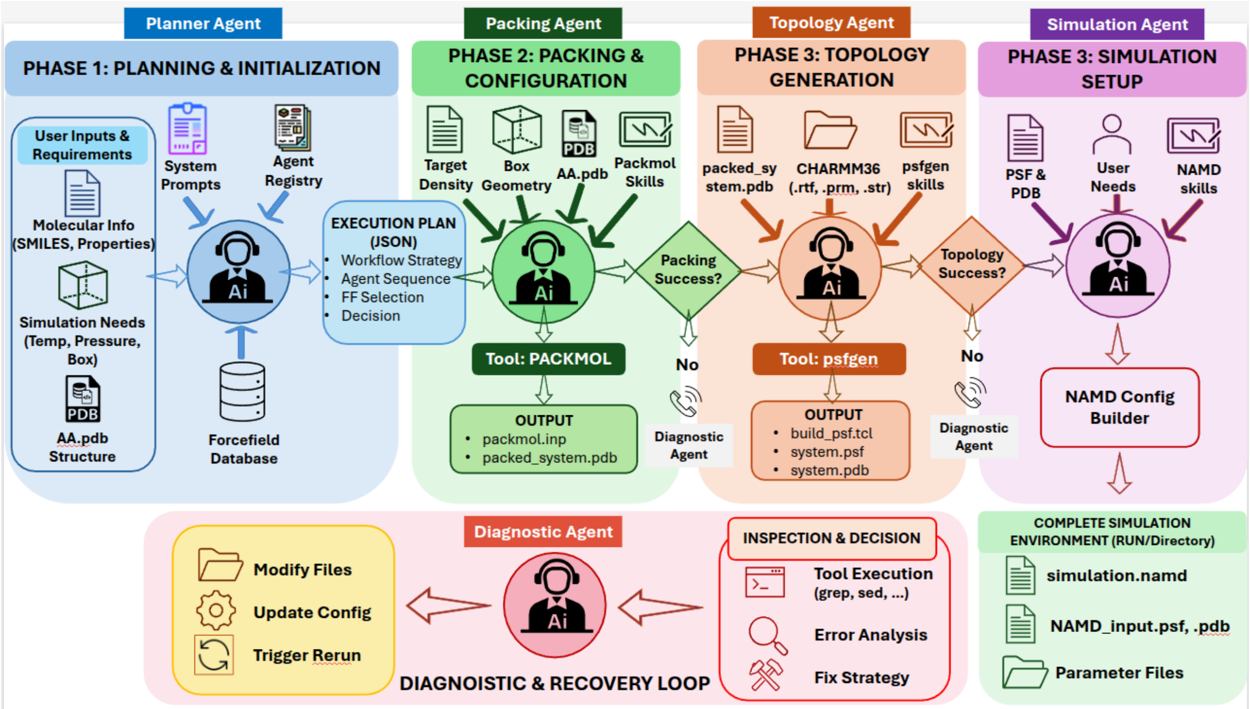


**Figure S1:** Schematic of the agentic AI orchestration for the all-atom equilibrium MD simulation setup with five specialized agents.

The multi-agent orchestration begins with user-provided molecular information, such as SMILES strings, physical properties (e.g., target density and molecular weight), and simulation requirements (e.g., temperature, pressure, and box dimensions). The Planner Agent analyzes these inputs to classify the molecular system as either a small molecule, protein, or conjugate class and selects required force fields for simulation, such as CHARMM36 for general biomolecular simulations or its CHARMM General Force Field[2] (CGenFF) extension for drug-like small molecules and solvents. The planner agent generates an execution plan in JSON format, outlining decisions on tool requirements (e.g., "Packmol" for molecular packing and "psfgen" for topology generation) and an ordered sequence of subsequent agents. This ordered planning phase ensures logical consistency, such as enforcing periodic boundary conditions for bulk liquid phases and validating input file existence to preempt failures.

Following planning, the Packing Agent automates the initial configuration of the molecular system using Packmol, a tool designed for efficient packing of molecules into predefined volumes to achieve target densities. For NAMD-compatible setups, the process involves calculating the number of molecules required for the user-specified box size (e.g., a 50 Å cubic box) based on the target density, generating a Packmol input script with tolerance parameters (typically 1.5–2.5 Å), and executing the packing via subprocess calls. The output

PDB file with the packed molecules is validated against density deviations (threshold <1% for accuracy), ensuring a physically realistic starting configuration suitable for CHARMM36 force field application. Any failure in this step triggers DiagnosticAgent for iterative fixing of the Packmol input file or the PDB file by reading the failure reasons from the output file, using its chemical knowledge augmented by the Packmol-specific skillset.

Topology generation is handled by the Topology Agent, which employs psfgen, a plugin from the Visual Molecular Dynamics (VMD)[3] suite, to assign CHARMM36-compatible atom types, charges, bonds, angles, and dihedrals to the packed system. The agent loads relevant topology files (e.g., top_all36_cgenff.rtf) and parameters (e.g., par_all36_cgenff.prm), creates a TCL script for segment definition and coordinate guessing, and produces PSF and PDB files essential for NAMD input. Validation steps confirm atom count consistency and residue naming compatibility, with automatic aliasing for mismatches through the DiagnosticAgent. This step is critical for CHARMM36-based simulations, as the force field additive all-atom model relies on precise parameterization to reproduce experimental properties.

Diagnostic Agent is a critical part of the agentic workflow, which employs a three-phase process: inspection, decision, and recovery to handle failures autonomously. Upon detecting errors (e.g., from Packmol density mismatches or psfgen parameter conflicts), it plans tool calls from a registry (e.g., grep for error patterns or sed for file modifications), analyzes results to categorize issues, and applies targeted fixes before retrying the failed agent. This robust error-handling mechanism, with specified retry budgets and confidence scoring, enhances workflow reliability in all-atom simulation setup.

At last, the Simulation Agent finalizes the NAMD environment by generating a comprehensive configuration file tailored to the user's specifications, such as an NPT ensemble at 298 K and 1 atm for a 100 ns production run with 2 fs timestep. The agent reads the example config files and skill set provided, which enables it for error-free one-shot config file creation. Moreover, it incorporates all required CHARMM36 parameters, sets periodic boundary conditions via cell basis vectors, and includes output controls for trajectories and restarts. For advanced setups, it supports extensions like steered MD or umbrella sampling, copying all necessary files into a dedicated “Run” directory before starting the simulation on HPC. This agentic integration ensures reproducibility and scalability, reducing setup time from days to hours while maintaining fidelity to established MD protocols.

**All-Atom Simulation Results:**

**Table S1:** The table lists the equilibrated thermodynamical properties at 298 K calculated on the last 30 ns out of 100 ns all-atom MD trajectory. The percentage error is calculated based on the deviation from the experimental values and reported within the braces.

| Solvent | Density ($g/cm^3$) | Heat of Vaporization (kcal/mol) | Surface Tension (dyn/cm) |
|---|---|---|---|
| DMSO | 1.0886 (0.58%) | 13.51 (6.82%) | 44.36 (5.39%) |
| DMA | 0.9448 (0.94%) | 13.98 (27.66%) | 42.58 (31.30%) |

After obtaining the atomistic trajectory from the bulk equilibrium simulation, it is converted into a CG trajectory using the coarse-grained mapping scheme proposed by the CG-MappingAgent. The equilibrium bonded parameters (bonds, angles, and dihedrals) and their fluctuations and the Rmin values from the RDF peaks are calculated and passed to the CG-BoundaryAgent for informed coarse-grained parameter boundary choices.

**Coarse-Grained (CG) Simulation Setup**

All molecular dynamics simulations were performed using NAMD (NAnoscale Molecular Dynamics)[1] with the CHARMM36[2] style force field. Three distinct simulation systems were prepared to calculate thermophysical properties:

Bulk liquid simulation: A cubic box containing N molecules with periodic boundary conditions (50 × 50 × 50 $Å^3$) was equilibrated under NPT conditions at 298K and 323K for DMSO and 298K and 313K for DMA, respectively. The barostat is kept at 1 atm pressure. The number of molecules in the simulation box is determined by the experimental density.

Liquid-vapor interface simulation: The final configuration from the bulk simulation was extended along the z-axis to create a slab geometry (150 Å) with vacuum regions above and below the liquid phase for surface tension calculations.

Gas-phase monomer simulation: A single molecule was simulated in a cubic box (50 × 50 × 50 $Å^3$) to determine the gas-phase energy.

All simulations are carried out with a 10 fs timestep with Langevin dynamics for temperature control (damping coefficient $\gamma = 1\ ps^{-1}$). Non-bonded interactions were calculated using a 12 Å cutoff with switching beginning at 9 Å. The bulk liquid simulation used a Parrinello-Rahaman barostat[4] to maintain constant pressure at 1 atm, while the interface and gas-phase simulations for surface tension calculation were performed at constant volume.

**Property Calculations**

The liquid density ρ was calculated from the equilibrated NPT simulation using the last 30 ns,

$$\rho = \frac{N \cdot Mw \cdot k}{\langle V \rangle}$$

where N is the number of molecules, $Mw$ = 78.14 g/mol and 87.12 g/mol are the DMSO and DMA molecular weights, k = 1.660578 g/(Å$^3$·amu) is the conversion factor, and ⟨V⟩ is the time-averaged volume extracted from the simulation output.

The molar heat of vaporization $\Delta H_{vap}$ was computed using the potential energy difference between the liquid and gas phases as,

$$\Delta H_{vap} = RT - \langle U_{liquid} \rangle / N + \langle U_{gas} \rangle$$

where R = 1.987 × 10$^{-3}$ kcal/(mol·K) is the gas constant, T = 298 K is the temperature, $\langle U_{liquid} \rangle$is the average potential energy of the liquid phase (averaged over the last 30 ns), N is the number of molecules, and $\langle U_{gas} \rangle$ is the average potential energy of a single molecule in the gas phase. The term RT = 0.5955 kcal/mol at 298 K accounts for the pV work.

The surface tension γ was calculated from the difference between the normal and tangential components of the pressure tensor in the slab simulation as,

$$\gamma = \frac{1}{2} L_z \left\langle P_{zz} - \frac{P_{xx} + P_{yy}}{2} \right\rangle$$

where $L_z$ = 150 Å is the box length in the z-direction, $P_{zz}$ is the normal pressure component, and $P_{xx}$ and $P_{yy}$ are the tangential components. The pressure components were converted from NAMD internal units (bar·Å) to dyn/cm using the conversion factor 0.01.

**Coarse-Grained Simulation Protocol During Agentic AI Optimization**

During the optimization loop, each simulation followed a two-stage protocol: (1) energy minimization for 1,000 steps using the conjugate gradient algorithm to remove unfavorable contacts, and (2) production runs of 20 ns. Coordinates and energies were saved every 100 steps for subsequent analysis.

**Coarse-Grained Agents' Workflow**

We used the open-source Moonshot AI's Kimi K2.5 LLM model (see model card on Hugging Face: https://huggingface.co/moonshotai/Kimi-K2.5) hosted on Virginia Tech ARC's computational cluster for the agentic workflow. The agent's communications were channeled through API requests, and the responses were logged into JSON and TXT-formatted files.

The **Mapping Agent (MA)** is responsible for proposing chemically plausible CG bead representations based on molecular topology and all-atom reference data. The agent reasons over atomic connectivity, functional groups, and charge distribution to generate multiple candidate mapping schemes while preserving molecular integrity. To prevent physically invalid groupings, the MA incorporates a connectivity-aware validation step that parses SMILES representations and bonding patterns, ensuring that only bonded atoms are grouped within a CG bead. For polar molecules, the agent is capable of introducing charged dummy beads to explicitly capture electrostatic interactions and preserve dipole orientation when required by molecular polarity.

The **Topology Agent (TA)** operates downstream of the MA and translates the selected CG mapping scheme into simulation-ready input files. The agent is provided with the CG mapping definition along with the all-atom PDB structure of the solvent molecule and generates topology and coordinate files (.psf and .pdb) compatible with the NAMD simulation package. To construct simulation boxes with user-specified dimensions, the TA leverages tool-calling capabilities to invoke external utilities, including Packmol and PSFgen, enabling automated generation of condensed-phase systems without manual intervention.

The **Boundary Agent (BA)** defines physically meaningful search boundaries for force-field parameter optimization. The correct selection of parameter bounds is critical for optimization efficiency, as overly restrictive bounds can bias the solution while excessively broad bounds can lead to unstable simulations. To address this, the BA is equipped with a domain-specific molecular dynamics skill set that enables geometry-based estimation of CG bead sizes and chemically motivated constraints on bonded and non-bonded interactions. When atomistic simulation data are available, the agent maps the all-atom trajectory onto the CG representation using the selected mapping scheme and extracts bond length, angle, and radial distribution function (RDF) statistics to estimate initial bounds based on the mean and standard deviation of the mapped distributions. In the absence of atomistic data, the BA estimates Lennard–Jones σ parameters from molecular geometry using radius-of-gyration calculations, with corrections applied for bond order, molecular flexibility, and electronic structure effects. Initial bounds for bond force constants, equilibrium bond lengths, and non-bonded ε and $R_{min}$ parameters are similarly derived using chemically informed heuristics. During optimization, these parameter boundaries are adaptively updated through expansion, contraction, and shifting operations based on diagnostic feedback, allowing the search space to evolve in response to optimization progress while maintaining physical consistency.

The parameter bounds are automatically optimized by the BA during optimization epochs. Specifically, parameter distributions from the top five performing solutions were analyzed using three primary boundary update operations: expansion, contraction, and shifting. Unilateral boundary expansion was triggered when parameter values clustered near a single boundary limit, indicated by at least two solutions lying within 10% of the minimum or maximum bound or when boundary hits exceeded two occurrences. Bilateral expansion occurred when solutions approached both bounds simultaneously, suggesting insufficient search space. Conversely, contraction was initiated when less than 25% of the parameter range was utilized and at least three solutions clustered in the interior region. The dynamical boundary change (see **Figure S7**) ensures a bounded, finite, yet enriched exploration space for the OA, which leads to faster convergence to the target properties.

The **Hypothesis Agent (HA)** generates physically plausible hypotheses for CG model parameters based on molecular reasoning and accumulated simulation history. At each iteration, the HA reads the performance scores across all parallel forks together with the diagnostic report produced by the Diagnostic Agent (DA) to formulate an updated hypothesis. This hypothesis is then passed to the Optimization Agent (OA), which translates it into concrete parameter sets for the next round of simulations. To evaluate each hypothesis, the HA coordinates parallel multi-fork simulations (nfork = 2, 4, or 8), wherein multiple candidate parameter sets are executed simultaneously. Higher nfork values broaden parameter space exploration and are expected to reduce the number of epochs required for convergence, at the cost of increased computational overhead per iteration. The aggregated simulation outcomes provide a statistically richer basis for parameter updates and enable increasingly accurate hypothesis generation in successive iterations.

The **Optimization Agent (OA)** proposes force-field parameter updates based on the current hypothesis from the HA, using adaptive multi-objective optimization strategies aimed at minimizing deviations between simulated and experimental target properties. The OA operates within the parameter bounds defined by the BA and incorporates feedback from the Diagnostic Agent (DA) to iteratively refine its proposals. To support stable convergence, the OA is equipped with three complementary memory types. A long-term memory stores historically best- and worst-performing parameter sets along with the associated LLM reasoning, enabling recall of effective parameter-update patterns and avoidance of unphysical regions of parameter space. A short-term memory records information from the most recent iterations to capture local optimization trends and mitigate oscillatory updates caused by noisy simulation feedback. An episodic memory records milestone states at fixed intervals to track macroscopic optimization progress and trigger higher-level adjustments to the optimization strategy when stagnation is detected.

The **Diagnostic Agent (DA)** analyzes simulation outputs generated during the optimization process and translates raw trajectory and thermodynamic data into structured feedback consumed by both the HA and OA. This includes evaluation of target thermodynamic properties, structural metrics, and phase behavior, as well as monitoring numerical instabilities, phase separation, or crystallization. Where parameter boundaries require revision, the DA communicates recommendations directly to the BA, closing the feedback loop across the three agents most central to iterative refinement. The DA thereby serves as the interpretive backbone of the closed-loop workflow, ensuring that subsequent hypothesis generation and parameter updates are grounded in a consistent and physically informed assessment of simulation outcomes.

Finally, a **Master Agent (MAS)** oversees the coordination of all six agents described above and manages the execution of the closed-loop workflow. The MAS orchestrates inter-agent communication, enforces execution order, monitors convergence criteria, and ensures stable and consistent operation of the autonomous optimization loop. Each agent operates under dedicated system prompts and user prompts, where system prompts define the agent's role, constraints, and reasoning logic, and user prompts provide task-specific context, feedback, and data required for execution.

**Fitness function used to drive the optimization loop**

The OA minimizes a composite error score that quantifies how well a given set of CG force-field parameters reproduces target experimental thermodynamic properties. At each optimization iteration, CG-MD simulations are run at two temperatures (298 K and 313 K for DMA; 298 K and 323 K for DMSO). For each temperature, the per-temperature score is computed as the sum of the percent absolute deviations of the simulated properties from their experimental target values. The composite score is then obtained by averaging the per temperature scores across all simulation temperatures, and it is this single scalar that the optimizer seeks to minimize. The properties included depend on schemes: Scheme 1 uses density ($\rho$), heat of vaporization (Hvap), and surface tension ($\gamma$), while schemes 2 and 3 additionally include the molecular dipole moment ($\mu$).

Per-temperature score is calculated as

$$E_T = 100 \times \sum \frac{\left| p_T^{expt} - p_T^{simulation} \right|}{p_T^{expt}}$$

where $p_T^{simulation}$ is the simulated value of property p at temperature T, and $p_T^{expt}$ is the corresponding experimental reference value.

Composite fitness score (minimization objective) is then calculated as

$$E = \frac{1}{N_T} \sum_{1}^{N_T} E_T$$

where $N_T$ is the number of simulation temperatures and the outer sum runs over all temperatures T.

**Agentic Prompts for the Coarse-Grained model development**

The agents are controlled by a two-tier prompt system designed to ensure reproducibility, scientific rigor, and adherence to strict physical and chemical constraints. The “system prompt” is persistent and directive: it establishes the agent as an expert in coarse-grained molecular modeling and force field development. This prompt enforces rigorous rules, such as the requirement for each core bead to contain at least two heavy atoms for the mapping agent and instructions on how to diagnose the system behavior by analyzing thermodynamic properties for the diagnostic agent. It also defines the mandatory output format as a valid JSON containing both reasoning and a detailed mapping schema. The “user prompt”, in contrast, is specific and iterative: it supplies the current molecule’s key properties and feedback on previous attempts, such as current results, validation errors, or crash information. This combination enables the agent to generate constraint-compliant, chemically meaningful workflow while allowing continuous refinement across iterations.

For example, the “system prompt” of the mapping agent includes

- Minimum of 2 heavy atoms (C, N, O, S, etc.) per core bead
- Strict connectivity rule: atoms within a bead must be either directly bonded or both bonded to the same heavy atom (1st-order neighbors)
- Grouping by chemically meaningful functional units rather than arbitrary connectivity

- Mandatory use of two dummy beads (DUP and DUN) when dipole moment is greater than 0.1 Debye, positioned on different core beads to correctly represent the molecular dipole vector
- No VdW interactions for dummy beads; only electrostatic interactions
- Symmetric interaction matrix for VdW parameterization (excluding dummies)
- Output restricted to valid JSON with predefined schema (reasoning + mapping)

This “system prompt” ensures that all generated mappings are physically meaningful, chemically accurate, and computationally efficient. On the contrary, the “user prompt” is instance-specific and provides molecule-specific information and feedback from previous iterations. It guides the agent toward improved mappings by incorporating:

- Molecule name, SMILES, 2D/3D connectivity structure
- Key physicochemical properties (polarity, dipole moment, molecular weight, total atom count)
- Molecular connectivity matrix and connected components
- Previously rejected mappings and their validation errors (to prevent repetition)
- Target properties to reproduce (e.g., density, heat of vaporization, surface tension, etc.)

The full prompts used for all agents are provided on GitHub.

**Skillset for the CG Agents:**

A comprehensive skillset is included in the Boundary Agent’s “system prompt” for the systematic estimation of coarse-grained (CG) force-field parameters, including bonded interactions, nonbonded Lennard-Jones terms, electrostatics, and effective bead sizes (σ). The skillset informs the agent about the loss of atomic degrees of freedom associated with coarse graining, and direct transfer of all-atom parameters is to be avoided. Instead, CG parameters are inferred by relating molecular geometry, bond topology, and interaction strengths to emergent structural, dynamical, and thermodynamic properties. Bonded interactions (bonds, angles, and dihedrals) are interpreted in terms of their roles in controlling molecular size, rigidity, conformational flexibility, and characteristic vibrational behavior, with force constants and equilibrium values softened to reflect the effective potentials required at the CG resolution. Nonbonded parameters are treated as collective interactions representing averaged dispersion, excluded volume, and cohesive effects, such that increases in Lennard-Jones well depth (ε) and bead diameter (σ or Rmin) are associated with higher density and viscosity and reduced diffusivity, while effective charges are used to encode screened electrostatic interactions and solvation behavior.

When atomistic mapping data are not available, CG bead sizes are estimated using a geometry-based procedure in which approximate three-dimensional atomic arrangements are reconstructed from bond lengths, bond orders, and angles, followed by the computation of the mass-weighted gyration radius of the mapped group. Physically motivated corrections accounting for van der Waals extent, bond flexibility, and electronic structure are subsequently applied. In this manner, the influence of molecular geometry (e.g., linear, bent, planar, or tetrahedral) and bond order on bead compactness is naturally captured, resulting in systematic variations in excluded volume and corresponding shifts in the peak positions of the radial distribution function. The estimated σ values are then converted into Lennard-Jones potential minima and CG bond lengths using consistent geometric relationships, ensuring internal force-field coherence. Overall, autonomous construction of chemically informed CG models is enabled, balancing structural fidelity, thermodynamic consistency, and computational efficiency while implicitly incorporating many-body, entropic, and averaging effects absent from direct all-atom parameter transfer.

**Parameter Guidance and Hypothesis Evolution:**

The question of whether an LLM-driven optimization agent engages in genuine physical reasoning or merely performs implicit parameter-space search is central to any rigorous evaluation of the value added by agentic frameworks relative to established stochastic and probabilistic optimization methods, including particle swarm optimization (PSO) and Bayesian optimization (BO). Residual coupling — defined as the rate at which a newly proposed hypothesis responds systematically to the dominant residual error of the preceding iteration — constitutes a necessary but insufficient criterion for distinguishing reasoned from reflexive optimization behavior: a naive gradient-following procedure would satisfy residual coupling by construction, in the absence of any underlying physical inference. The distinguishing behavioral signature of a reasoning agent lies instead in its propensity to invoke named physical laws, recognize and respect geometric constraints, diagnose the mechanistic origins of iterative failures, and formulate multi-parameter perturbations that explicitly account for inter-property coupling.

To address this question, we report three complementary quantitative metrics computed across seventeen Scheme 3 optimization runs (nine in DMSO, eight in DMA): a behavioral residual-coupling rate, a behavioral crash-response rate, and a linguistic mechanism-signature rate. The first two metrics are summarized by cohort in **Figures S3–S4**, with per-run breakdowns provided in **Figures S5–S6**. The third metric, which constitutes the primary quantitative basis for the mechanism-level reasoning analysis, is the focus of the corresponding main-text section and is summarized in **Figure S2**.

**Mechanism-level reasoning signatures**

We scanned the scientific with non-empty rationale text of every hypothesis in the Scheme 3 cohort and classified each rationale according to whether it contained one or more of six mechanism-level reasoning signatures defined by case-insensitive regular-expression patterns. The six signatures are non-exclusive; a single rationale frequently matches two or three of them.

| Signature | What it captures |
|---|---|
| **Compensation moves** | Hypothesis explicitly balances a change in one parameter with a counter-change in another to preserve a property. Keywords: compensate, maintain, preserve, while simultaneously, proportional, offset, balance. |
| **$\mu = q \times d$ invocation** | Hypothesis explicitly cites the dipole-charge-distance relationship. Matches ' $\mu = q \times d$', '$q \times d$', 'charge separation', and equivalent forms. |
| **Dummy-inside-core constraint** | Hypothesis explicitly names the geometric constraint that charged dummy beads must reside within their parent bead's LJ sigma/2 radius. Matches 'dummy inside core', 'sigma/2', 'must reside'. |
| **Crash root-cause** | Hypothesis diagnoses the mechanism behind a simulation crash using causal language. Matches 'because', 'due to', 'caused by', 'catastrophe', 'singularity', 'without VdW shielding'. |
| **Boundary-aware pivot** | Hypothesis recognizes that a parameter has reached a physical or numerical limit and explicitly switches to a different physical lever. Matches 'at maximum allowed', 'cannot reduce further', 'fixed at boundary', 'at upper/lower bound'. |
| **Coupled trade-off** | Hypothesis explicitly discusses coupling between target properties. Matches 'electrostatic component of', 'opposing effects', 'competing requirements', 'trade-off', 'without exacerbating', 'coupled'. |

Across the seventeen-run Scheme 3 cohort, a total of 642 hypotheses scientific rationale were scanned. Of these, 467 (72.7%) contained at least one signature. Per-signature counts are

| Signature | Hits | Fraction |
|---|---|---|
| **Compensation moves** | 245 | 38.2% |
| **$\mu = q*d$ invocation** | 237 | 36.9% |
| **Dummy-inside-core** | 101 | 15.7% |
| **Crash root-cause** | 84 | 13.1% |

| | | |
|---|---|---|
| **Boundary-aware pivot** | 66 | 10.3% |
| **Coupled trade-off** | 42 | 6.5% |
| **At least one signature** | 467 | 72.7% |

Breakdown by solvent for scheme 3

| Signature | DMSO (n=381) | DMA (n=261) |
|---|---|---|
| **Compensation moves** | 35.4% | 42.1% |
| **μ = q*d invocation** | 25.7% | 53.3% |
| **Dummy-inside-core** | 16.8% | 14.2% |
| **Crash root-cause** | 14.4% | 11.1% |
| **Boundary-aware pivot** | 7.9% | 13.8% |
| **Coupled trade-off** | 7.9% | 4.6% |
| **At least one signature** | 68.0% | 79.7% |

The signature hit rates across all six cases

| Cohort | Any | Compensation | μ = q*d | Dummy-inside-core | Crash | Boundary-aware | Coupled |
|---|---|---|---|---|---|---|---|
| **DMSO Scheme 1** | 60.4% | 41.7% | 0.3% | 0.0% | 17.5% | 6.3% | 12.4% |
| **DMSO Scheme 2** | 72.0% | 35.3% | 32.4% | 14.6% | 13.6% | 12.4% | 7.3% |
| **DMSO Scheme 3** | 68.0% | 35.4% | 25.7% | 16.8% | 14.4% | 7.9% | 7.9% |
| **DMA Scheme 1** | 54.2% | 39.0% | 0.9% | 0.9% | 14.3% | 6.7% | 9.9% |
| **DMA Scheme 2** | 63.0% | 35.7% | 19.4% | 8.2% | 13.5% | 4.4% | 13.5% |
| **DMA Scheme 3** | 79.7% | 42.1% | 53.3% | 14.2% | 11.1% | 13.8% | 4.6% |

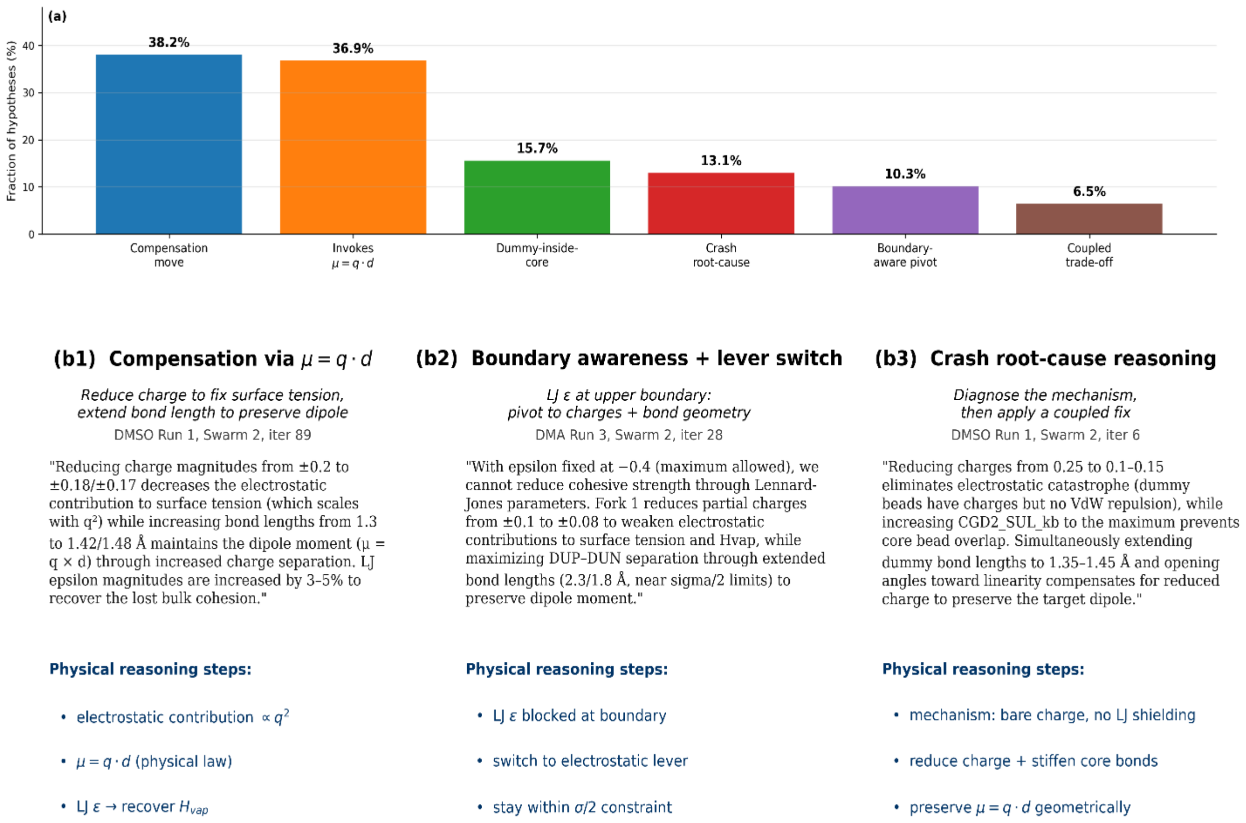


**Figure S2:** (a) Fraction of the 642 hypotheses in the seventeen-run Scheme 3 cohort that contains each of the six mechanism-level reasoning signatures. 72.7% of hypotheses contain at least one signature. (b1) Compensation via μ = q × d: the agent reduces dummy-bead charge to lower the electrostatic contribution to surface tension and extends the dummy-bead bond length to preserve the dipole moment. (b2) Boundary-aware pivot: the agent recognizes that LJ ε has reached its maximum allowed value and switches to the electrostatic and bond-geometry levers. (b3) Crash root-cause reasoning: the agent diagnoses the physical mechanism behind a velocity-overflow crash (bare charges on dummy beads without van der Waals shielding) before proposing a coupled fix.

**Linked subsystem to parameters**

For Scheme 3, 16 parameters for DMSO and 28 parameters for DMA could be assigned, a priori, to exactly one of five physical subsystems (LJ nonbonded, Electrostatics, Bond geometry, Bond/angle stiffness, Angle geometry). The assignment reflects which physical quantity each parameter controls in a CG force field and is fixed once per Scheme before any metrics are computed. Each of the four target properties is then linked to the subset of these five subsystems that can physically move it. These linkages are not chosen after inspecting the results — they come from the textbook physics of the CG force field, in which

LJ parameters set bulk cohesion and packing (controlling density, heat of vaporization, and surface tension), while the molecular dipole moment is determined by the dummy-bead charge magnitudes and the dummy-to-core bond lengths through the relationship $\mu = q \times d$. The resulting target-to-subsystem mapping is:

| Target property | Linked subsystems | Physical justification |
|---|---|---|
| **Density** | LJ nonbonded; Bond geometry | Excluded volume is set by LJ $R_{min}$; intermolecular cohesion by LJ ε; bond length sets molecular extent and packing. |
| **Heat of vaporization** | LJ nonbonded; Electrostatics | Cohesive energy is dominated by LJ ε plus the electrostatic component contributed by the charged dummy beads. |
| **Surface tension** | LJ nonbonded | At CG resolution, interfacial cohesive forces are almost entirely carried by LJ ε; other subsystems contribute negligibly. |
| **Dipole moment** | Electrostatics; Bond geometry | $\mu = q \times d$, where q is the dummy-bead charge magnitude (Electrostatics) and d is the dummy-to-core separation set by bond geometry. |

An event is classified as a 'parameter hit' or equivalently when the hypothesis is said to perturb a linked subsystem, if the parameters being changed list of that hypothesis contains at least one parameter belonging to any of the subsystems linked to the current dominant residual. For example, if the dominant residual at iteration $i - 1$ is the dipole moment, and the hypothesis at iteration $i$ proposes to change {DUP_charge, DUN_charge, CGD2_DUP_bl, SUL_epsilon}, then two of the four perturbed parameters (DUP_charge, DUN_charge) belong to Electrostatics and one (CGD2_DUP_bl) belongs to Bond geometry, which are both linked to dipole moment; the event is therefore a parameter hit regardless of the fourth parameter (SUL_epsilon is in LJ nonbonded, which is not linked to dipole). DUP and DUN denote the dummy positive and negative beads. A single perturbed parameter in a linked subsystem is sufficient for the event to count as a hit.

The parameter-hit criterion is stricter than it may appear. Under a uniform-random hypothesis policy that draws parameters without regard to the dominant residual, the probability of hitting a linked subsystem is roughly the linked subsystem's share of the parameter count which is approximately 30-50% depending on the property. Observed parameter-hit rates of 83-97% (panel (a) of Figures **S3** and **S4**) therefore rule out random search at statistical confidence and constitute the behavioral half of the residual-coupling test.

**Residual coupling and crash response**

For each iteration *i > 1* with a non-empty hypothesis, we computed the normalized residual errors at iteration *i − 1* and identified the dominant residual as the property with the largest absolute percent deviation from its experimental value. An iteration was scored as responding to the dominant residual if the hypothesis at *i* either mentioned that property in its scientific rationale text or perturbed at least one parameter in a physically linked subsystem.

To test whether the hypothesis at iteration *i* responded to that dominant residual, using two independent criteria. The first criterion 'text hit' (blue bars in panel (a) of **Figures S3** and **S4**) is satisfied when the hypothesis text mentions the dominant property by name, using a case-insensitive regular-expression match against a small property-specific lexicon. *For density: 'density', 'packing', 'excluded volume', 'rmin', 'sigma', 'rminby2'. For heat of vaporization: 'heat of vap', 'Hvap', 'vaporization', 'cohesion', 'epsilon'. For surface tension: 'surface tension', 'interfacial', 'gamma'. For dipole moment: 'dipole', 'charge', 'polarity', 'dummy'*. The second criterion 'parameter hit' (orange bars) is satisfied when the hypothesis perturbs at least one parameter in a subsystem linked to the dominant property, as defined in the preceding subsection.

Panel (b) measures how the HA responds to simulation failures rather than to property-error feedback. For every iteration whose diagnostic warnings contained at least one crash marker (*regex match for 'fatal', 'instability', 'velocity overflow', 'singularity', 'too fast', 'too small', or 'extreme force'*), we examined the hypothesis produced at the immediately following iteration of crash. A crash-responsive hypothesis is defined analogously to a residual-coupling hit, with two independent criteria:

(1) Text response: the next hypothesis contained both a reduction verb (*reduce, lower, soften, decrease, mitigate, prevent, avoid, stabilize, eliminate*) and a crash keyword, indicating that the agent explicitly acknowledged the failure mode in its own words.

(2) Parameter response: the next hypothesis perturbed at least one parameter in a subsystem physically linked to the classified crash cause. Charge singularities are linked to Electrostatics and Bond geometry (because reducing charge magnitude or extending dummy-bead bond lengths alleviates unshielded Coulombic forces); stiffness overflow is linked to Bond/angle stiffness (softening force constants damps high-frequency vibrations); steric clash is linked to LJ nonbonded and Bond geometry (expanding σ or extending bond lengths removes overlapping volumes); generic instabilities are permitted to respond via any of the four bonded or electrostatic subsystems. Panel (b) of **Figures S3** and **S4** reports these two rates, along with the 'either' rate, broken down by fork count (Fork 2, Fork 4, Fork 8) for DMSO and DMA, respectively.

Panel (c) summarizes where in parameter space the HA is spending its effort. At each iteration, we computed the fraction of perturbed parameters belonging to each of the five physical subsystems into which the Scheme 3 force-field parameters were grouped a priori:

| Subsystem | Physical meaning | Parameters (DMSO / DMA) |
|---|---|---|
| **LJ nonbonded** | ε and Rmin; control excluded volume, cohesion, and packing | 4 / 10 |
| **Electrostatics** | dummy-bead charges; control μ via charge magnitude | 2 / 2 |
| **Bond geometry** | bond lengths; control molecular extent and dummy separation | 3 / 4 |
| **Bond/angle stiffness** | force constants; control vibrational modes and stability | 5 / 8 |
| **Angle geometry** | equilibrium angles; control molecular shape | 2 / 4 |

For each run we first averaged the per-iteration subsystem shares over all iterations in that run, then averaged the resulting run-level shares across the replicates at each fork count. The resulting 3 × 5 matrix is displayed as a heatmap in panel (c) of **Figures S3** and **S4**, with numerical values overlaid.

Pooled rates (event-level weighting):

| Metric | n events | DMSO rate | DMA rate | Pooled |
|---|---|---|---|---|
| **Residual coupling (either)** | 610 | 98.6% | 96.3% | 97.7% |
| **Crash response (either)** | 487 | 93.3% | 91.1% | 92.4% |

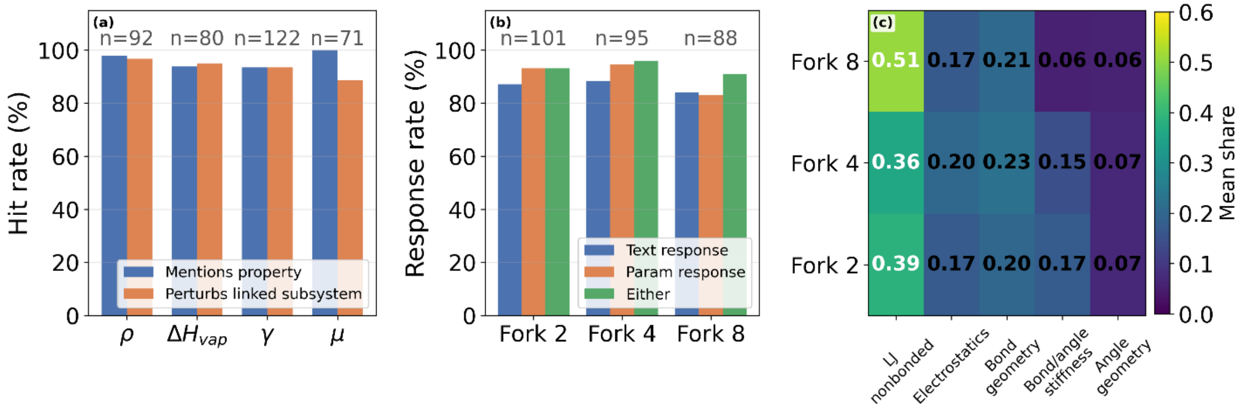


**Figure S3:** DMSO scheme-3 cohort (9 runs). (a) Residual-coupling hit rates are broken down by which property was the dominant residual. (b) Crash-response rates by fork count. (c) Mean share of perturbed parameters in each physical subsystem, averaged across replicates and iterations.

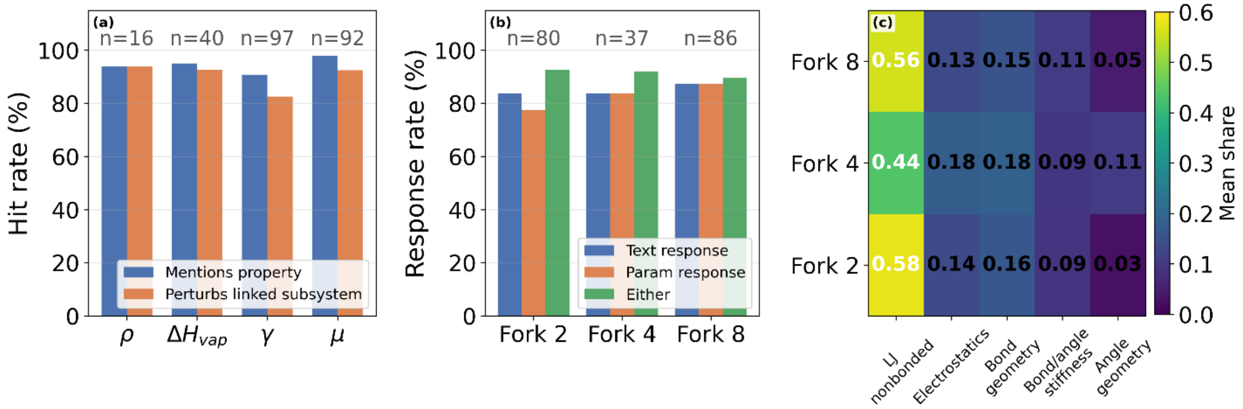


**Figure S4:** DMA scheme-3 cohort (8 runs). Same panel structure as **Figure S3**; the close agreement between the two figures shows that the reasoning behavior replicates across chemically distinct target solvents and across different coarse-grained topologies (DMSO: 16 parameters; DMA: 28 parameters).

Both DMSO and DMA, the LJ-nonbonded subsystem carries the largest share of perturbed parameters across every fork count, which is the physically expected behavior: LJ ε and $R_{min}$ are the primary control variables for three of the four target properties (density, heat of vaporization, and surface tension), and only the dipole moment depends principally on the electrostatic and bond-geometry subsystems. Moreover, despite the scheme topologies being different (DMSO uses 2 core beads + 2 dummy beads at 16 total parameters; DMA uses 3 core beads + 2 dummy beads at 28 total parameters), the rank ordering of the subsystems is qualitatively identical: LJ nonbonded > Bond geometry ≈ Electrostatics > Bond/angle stiffness > Angle geometry (see panel (c)). The agent is therefore allocating its effort in a manner that respects the physical importance of each subsystem rather than the relative size of each subsystem — it is not simply perturbing whichever parameters happen to be most numerous.

For schemes 1 and 2, we observed a similar trend as for Scheme 3. Mean share of perturbed parameters in each subsystem, by cohort and by fork count. Scheme 1 has only three populated subsystems (LJ nonbonded, bond geometry, and bond/angle stiffness); the electrostatics and angle geometry columns are zero by construction because those subsystems contain no parameters in Scheme 1. All subsystems from three schemes are the following:

| Cohort | Fork count | LJ nonbonded | Electrostatics | Bond geometry | Bond/angle stiffness | Angle geometry |
|---|---|---|---|---|---|---|
| **DMSO Scheme 1** | Fork 2 | 91.4% | 0.0% | 4.9% | 3.7% | 0.0% |
| **DMSO Scheme 1** | Fork 4 | 73.0% | 0.0% | 16.3% | 10.7% | 0.0% |
| **DMSO Scheme 2** | Fork 2 | 39.6% | 16.6% | 20.7% | 11.7% | 11.4% |
| **DMSO Scheme 2** | Fork 4 | 36.5% | 22.2% | 21.8% | 10.1% | 9.5% |
| **DMSO Scheme 2** | Fork 8 | 36.2% | 19.2% | 21.1% | 12.5% | 10.9% |
| **DMSO Scheme 3** | Fork 2 | 38.6% | 17.3% | 19.9% | 17.4% | 6.7% |
| **DMSO Scheme 3** | Fork 4 | 35.7% | 20.2% | 22.6% | 14.6% | 6.9% |
| **DMSO Scheme 3** | Fork 8 | 50.7% | 17.0% | 20.6% | 5.6% | 6.0% |
| **DMA Scheme 1** | Fork 2 | 80.1% | 0.0% | 11.8% | 8.1% | 0.0% |
| **DMA Scheme 1** | Fork 4 | 82.6% | 0.0% | 10.4% | 7.0% | 0.0% |
| **DMA Scheme 1** | Fork 8 | 83.5% | 0.0% | 7.7% | 8.8% | 0.0% |
| **DMA Scheme 2** | Fork 2 | 53.0% | 17.8% | 19.2% | 3.6% | 6.4% |
| **DMA Scheme 2** | Fork 4 | 61.1% | 14.0% | 15.1% | 5.1% | 4.7% |
| **DMA Scheme 2** | Fork 8 | 53.4% | 14.9% | 17.1% | 9.4% | 5.1% |
| **DMA Scheme 3** | Fork 2 | 58.1% | 13.8% | 16.4% | 8.9% | 2.8% |
| **DMA Scheme 3** | Fork 4 | 43.6% | 17.7% | 18.5% | 9.4% | 10.7% |
| **DMA Scheme 3** | Fork 8 | 55.9% | 13.2% | 15.3% | 10.7% | 4.9% |

The residual coupling generalizes all three schemes for both solvents. All six cohorts lie in the 90–100% 'either' coupling band, and the per-property breakdown shows that each active property is coupled at ≥90% across every cohort in which it is active. Moreover, crash response generalizes as well across the four dummy-bearing cohorts (schemes 2 and 3 in both solvents), where the response rate stays within an 88–94% band. Scheme 1 produces substantially fewer crashes overall because the dominant failure mode (charge-singularity instabilities between oppositely charged dummy beads) cannot occur without dummy charges. The Scheme 1 crash response rates are lower (65% DMSO, 80% DMA), but the result reflects a metric-definition limitation. The crash-cause classifier maps Scheme 1 failures to a 'linked subsystem' (electrostatics) that is empty in Scheme 1, so the parameter-hit test under-credits physically valid LJ and bond-stiffness responses. The dummy-bearing cohorts provide a better test of the HA's crash-handling behavior for schemes 2 and 3.

Mechanism signatures generalize for the four non-electrostatic signatures (compensation moves, crash root-cause reasoning, boundary-aware pivots, and coupled trade-offs). LJ nonbonded is the most-perturbed subsystem in every cohort and fork count combination, with mean shares ranging from 35.7% (DMSO Scheme 3, Fork 4) to 73% in some Scheme 1 cohorts. A higher rate is observed for Scheme 1 as the linked subsystems related to electrostatic interactions are limited in the absence of dummy beads.

**Per-run breakdowns**

The aggregate rates are averaged over fork counts and replicates in Scheme 3. The per-run breakdowns in **Figures S5** and **S6** show that the response rates are stable across individual runs despite variation in iteration budgets, that the crash-cause distribution is dominated by charge-singularity warnings in both solvents, and that the LJ nonbonded subsystem retains the largest share of perturbed parameters throughout every phase of every run.

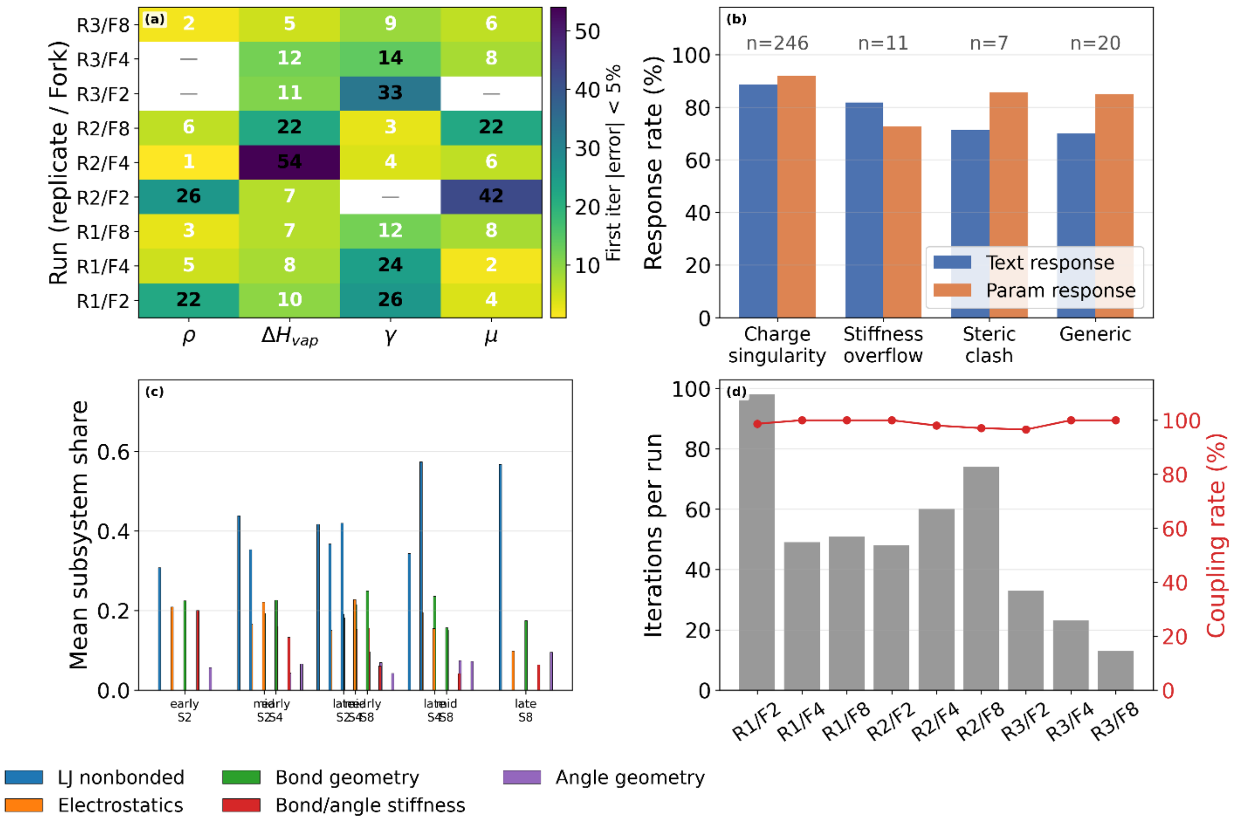


**Figure S5:** DMSO per-run breakdowns. (a) Iteration at which each property first entered ±5% of the experiment, per run. (b) Crash-response rates are broken down by crash-cause class. (c) Mean subsystems share by iteration phase (early/mid/late) and fork count. (d) Per-run iteration budget (grey bars) overlaid with the residual-coupling rate (red line).

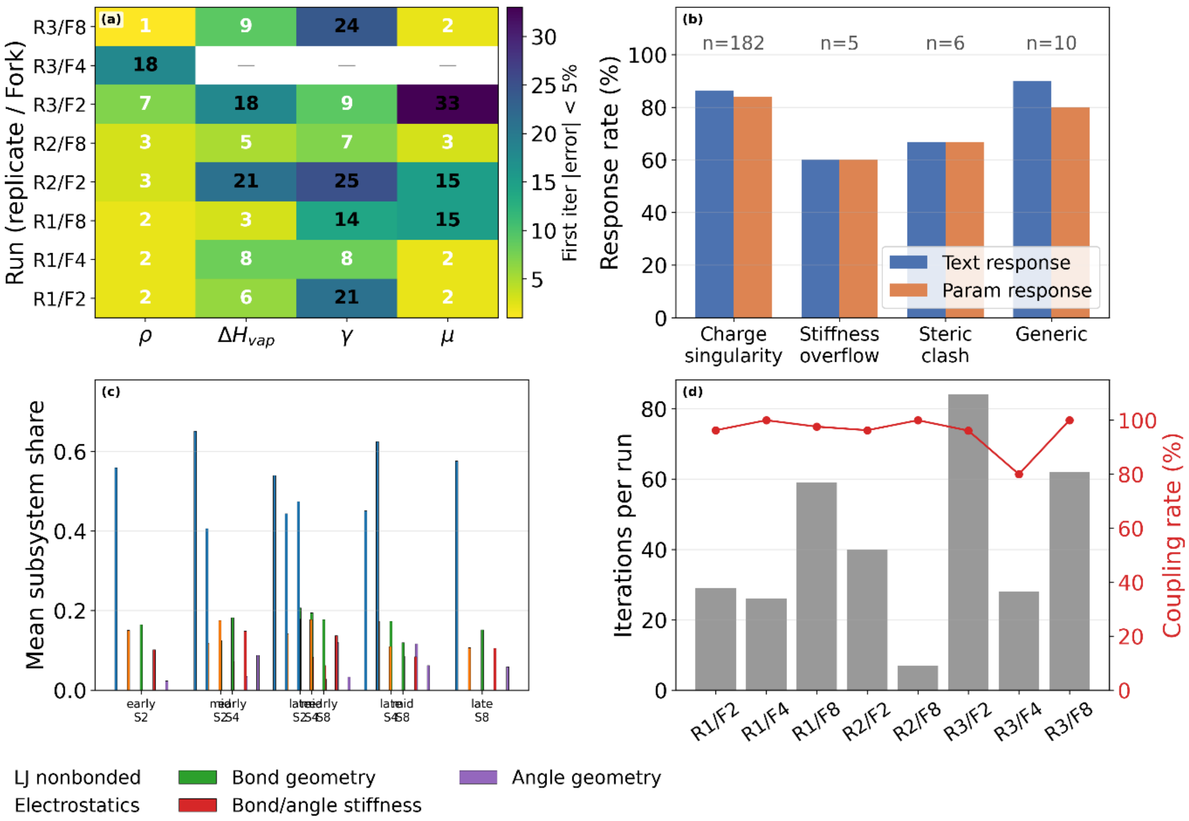


**Figure S6:** DMA per-run breakdowns. Same panel structure as **Figure S5**.

**Case study 1 — Compensation via μ = q × d (DMSO)**

This specific example is taken from DMSO Run 1, Fork 2, Scheme 3, epoch 89. The surface tension residual is dominant and overestimated; density and heat of vaporization are both slightly below target; dipole moment is close to target.

Rationale (verbatim):

> *Reducing charge magnitudes from ±0.2 to ±0.18/±0.17 decreases the electrostatic contribution to surface tension (which scales with $q^2$) while increasing bond lengths from 1.3 to 1.42/1.48 Å maintains the dipole moment (μ = q × d) through increased charge separation. To compensate for the reduced electrostatic cohesion in bulk properties, LJ epsilon magnitudes are increased by 3–5% toward their upper bounds.*

The agent simultaneously performs three mechanism-level reasoning steps. First, it identifies the electrostatic contribution to surface tension as scaling with $q^2$, invoking the quadratic dependence of Coulombic interactions on charge magnitude. Second, it recognizes that simply reducing charges would regress the dipole moment and cites μ = q × d by name to justify extending bond lengths as a compensating move that preserves the product. Third, it anticipates that reducing q will also reduce bulk cohesive energy (and therefore density and Hvap) and proposes a small LJ ε increase to recover the lost cohesion through a different physical mechanism. A single hypothesis thus contains three coupled moves, each justified by a distinct physical relationship. This case study is reproduced in **Figure S2 panel (b1)**.

**Case study 2 — Boundary-aware pivot (DMA)**

This specific example is taken from DMA Run 3, Fork 2, Scheme 3, epoch 28. Density is roughly 15% below target, Hvap is 15% above target, surface tension is 47% above target, dipole moment is dramatically below target. LJ ε has previously been pushed to its physical upper boundary of −0.4 in an attempt to increase cohesion.

Rationale (verbatim):

> *With epsilon fixed at −0.4 (maximum allowed), we cannot reduce cohesive strength to fix the 15% Hvap excess and 47% surface tension overestimation through Lennard-Jones parameters. Instead, Fork 1 reduces partial charges from ±0.1 to ±0.08 to weaken electrostatic contributions to surface tension and Hvap, while maximizing DUP–DUN separation through extended bond lengths (2.3/1.8 Å, near sigma/2 limits) and linearized angles (168°) to preserve dipole moment.*

The agent explicitly identifies that the LJ subsystem is no longer a usable lever because ε has reached its maximum allowed value. Rather than continuing to push against the boundary, it pivots to the electrostatic subsystem on the grounds that electrostatic forces also contribute to surface tension and heat of vaporization — a subtle physical claim about which subsystems contribute to which properties. Simultaneously it recognizes that reducing charges will regress the dipole moment and counters this by maximizing the DUP–DUN separation through extended bond lengths — staying within the sigma/2 geometric constraint identified separately. The parameter move is a textbook example of physical-constraint-aware multi-lever reasoning. This case study is reproduced in **Figure S2 panel (b2).**

**Case study 3 — Crash root-cause reasoning (DMSO)**

This specific example is taken from DMSO Run 1, Fork 2, Scheme 3, epoch 6. The previous iterations have produced velocity-overflow crashes with atoms exceeding the 1200 Å/ps velocity limit, flagged by the Diagnostic Agent as electrostatic instability.

Rationale (verbatim):

*Reducing charges from 0.25 to 0.1–0.15 eliminates electrostatic catastrophe (dummy beads have charges but no VdW repulsion), while increasing CGD2_SUL_kb to the maximum 20.0 prevents core bead overlap and LJ singularities. Simultaneously, increasing dummy bond lengths to 1.35–1.45 Å and optimizing angles toward linearity (175°/170° at CGD2, 125°/120° at SUL) maximizes DUP–DUN separation distance to compensate for reduced charge magnitude, attempting to preserve the target dipole moment of 3.96 D.*

The hypothesis begins with a mechanistic diagnosis: the crashes are caused specifically by charged dummy beads that carry no van der Waals repulsion of their own and therefore produce unshielded Coulombic forces when they approach closely. This is not merely 'reduce the thing that caused the crash'; it is an explanation of why reducing charges is the right fix, grounded in the physical structure of the coarse-grained representation. The hypothesis then constructs a multi-parameter response: reduce charges (primary fix), stiffen the core bond to prevent associated core-core overlap, and compensate for the lost dipole by extending dummy-bead bond lengths and linearizing angles. The fix is physically coupled, not a one-dimensional retreat from the failing parameter. This case study is reported in **Fig. S2 panel (b3)**.

**An illustrative example for a single run of DMSO**

As an illustrative single-run walk-through, we analyzed the hypothesis-driven optimization carried out by the Hypothesis Agent for DMSO Scheme 2 at Fork 4 (one of the 54 runs in our campaign). In this trajectory, the composite objective decreased from 182.85 at iteration 1 to 6.90 at iteration 49, a 96.22% reduction. Within this single run the hypothesis content can be organized into three informal regimes — exploratory, compensatory, and refinement — described in the paragraphs below. We emphasize that this regime structure is specific to the dynamics of this one run and should not be interpreted as a general temporal pattern: the cross-run mechanism-signature analysis in the preceding subsection (n = 642 hypotheses across 17 runs) shows that the Lennard-Jones nonbonded subsystem retains the largest share of perturbed parameters throughout every phase of every run and that no single temporal ordering replicates across the cohort. The single-run narrative below is therefore included as a qualitative walk-through of one trajectory rather than as evidence for a general three-phase structure.

**Exploratory Regime:** In the initial phase (iterations 1 to 6) the hypotheses were generated to perform high-dimensional exploration, averaging 8 parameter updates per iteration across 4 different parameter families. All major interaction classes are perturbed in every iteration (LJ ε, $R_{min}$, charges, and bond lengths: 5/5 occurrences each), yielding a MAPE of 22% across the four target properties in both temperatures. Proposed updates explicitly probe physically interpretable axes: cohesion (ε), excluded volume ($R_{min}$), and polarity (charges and bond geometry) under imposed constraints (e.g., charge and bonded parameter bounds), indicating structured exploration rather than stochastic sampling.

**Compensatory Regime:** By iteration 8, the optimizer resolves competing physical trends, reducing the MAPE to 4%. Diagnostic signals reveal underpredicted density (excessive excluded volume) coexisting with overpredicted heat of vaporization and surface tension (excessive cohesion). The HA responds with coordinated updates: "decreasing $R_{min}$ to improve packing, increasing ε (less negative) to reduce cohesion, and modestly reducing charge magnitudes". Multiple concurrent proposals systematically sample this compensation space, reflecting hypothesis-driven correction of coupled thermodynamic observables.

**Refinement Regime:** Near convergence (iterations = [28,37,40,44,49]), the search contracts to 5.6 parameter updates across 3.2 average families. LJ size is no longer modified (0/5 iterations), and LJ ε updates decrease (2/5 iterations), while electrostatic and bonded parameters dominate (charges, bond lengths, and stiffness: 4/5 iterations each). Notably, 3/5 iterations involve no LJ updates, indicating effective convergence of nonbonded terms. The MAPE in this regime is 6.69%, consistent with localized refinement. This phase also incorporates explicit stability corrections: reduction of bond stiffness, elongation of dipole-defining bonds, and softening of angular force constants to mitigate high-frequency instabilities. These updates demonstrate coupling between accuracy-driven and stability-driven objectives.

**Final State:** The optimal configuration (iteration 49) achieves a composite score of 6.90 and MAPE of 1.73%. Residual improvements are governed by targeted electrostatic and bonded adjustments, particularly dipole tuning via bond length modulation and stabilization via reduced force constants.

Overall, this single optimization follows a structured progression from global, high-dimensional exploration to low-dimensional, physically localized refinement by hypothesis generated with chemical reasoning coupled with feedbacks from DA. High-performing iterations involve systematically fewer parameter updates and parameter families than low-performing iterations, indicating that convergence is achieved through progressive constraint and physical interpretability, rather than continued broad exploration or stochastic search.

**Temperature -specific response of HA**

The optimization in this work is run simultaneously at two temperatures per solvent (DMSO: 298 K / 323 K; DMA: 298 K / 313 K). The HA receives DA reports that describe residual errors at both temperatures, and the fitness function penalizes parameter sets that fit one state point at the expense of the other. This extra constraint is a substantive increase in problem difficulty: a naive parameter sweep that optimizes a single-temperature metric will not generally pass a two-temperature acceptance test. The cross-solvent cross-scheme analysis in this document tests two propositions: first, that the HA explicitly reasons about the two-temperature fitness structure rather than treating the optimization as a single-state-point fit; and second, that the resulting parameter sets achieve comparable accuracy at both temperatures.

We scan every hypothesis in all six cohorts for six temperature language markers.

(a) Specific T numeric — the rationale contains the literal string '298K', '313K', or '323K'. This is the most direct textual evidence that the HA is citing a specific temperature.

(b) Temperature dependence (strict) — the rationale contains 'temperature' adjacent to one of 'dependen', 'sensitiv', 'transferab', 'scal', 'gradient', or 'derivative'. The earlier version also accepted 'range', 'trend', and 'response', which admitted too many false positives; these are now excluded.

(c) One-T-only mismatch — the rationale contains a qualifier ('only', 'while', 'but', 'whereas', 'however', 'although') followed within 60 characters by a numerical temperature token ('298 K', '313 K', '323 K'). The proximity constraint is new in this version and eliminates cases where 'only' and the temperature token were matched across arbitrary intervening text.

(d) Temperature trade-off (new marker) — the rationale explicitly contrasts behavior at two temperatures, using patterns such as 'at X K... while... at Y K', 'improve at X K... but worsen at Y K', 'at one temperature... at the other temperature', or the physical phrase 'too steep temperature dependence'. This marker is the cleanest indicator that the HA is reasoning about the two-temperature mismatch structure rather than about temperatures in isolation.

(e) Across-both phrasing — 'both temperatures', 'both state points', 'at both', 'across both', 'each temperature'.

(f) Anomalous temperature dependence — the rationale explicitly flags that a property's temperature response is wrong (e.g. 'decreases with T when it should increase', 'temperature slope too steep').

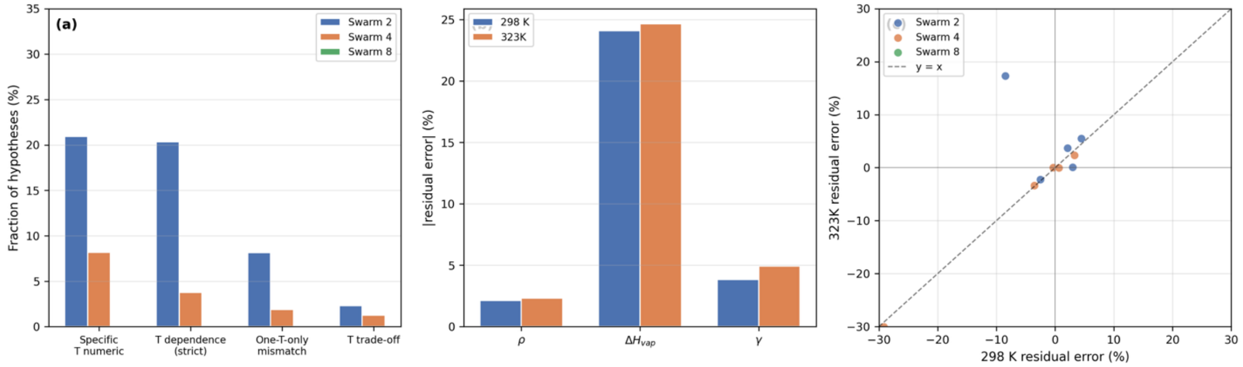


**Figure S7:** (a) Percentage of hypothesis that contained different temperature language markers across for DMSO Scheme 1 and forks/swarms (b) Residual error across different properties at two different temperatures (c) The residual error correlation between two temperatures.

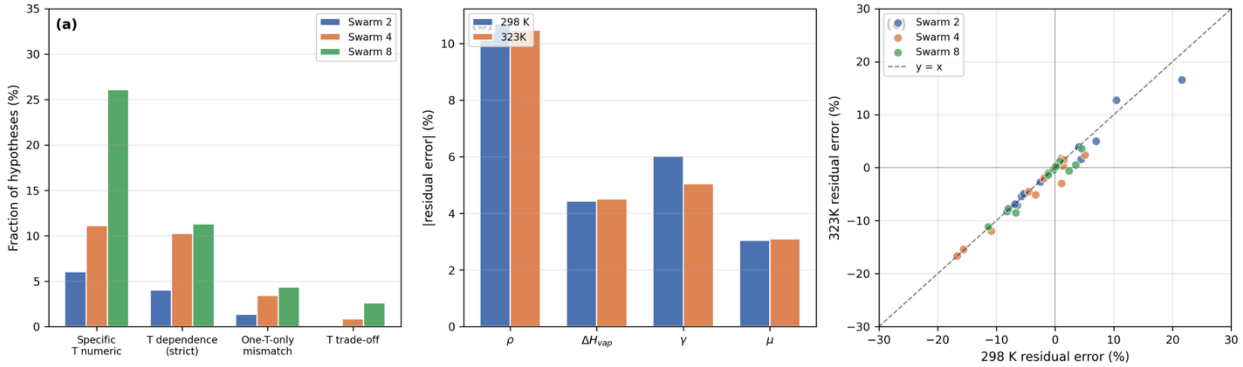


**Figure S8:** (a) Percentage of hypothesis that contained different temperature language markers across for DMSO Scheme 2 and forks/swarms (b) Residual error across different properties at two different temperatures (c) The residual error correlation between two temperatures.

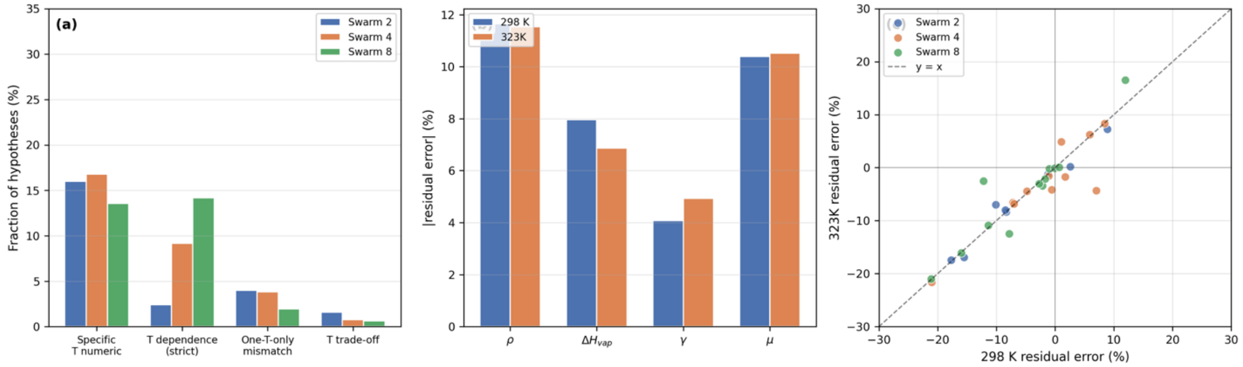

**Figure S9:** (a) Percentage of hypothesis that contained different temperature language markers across for DMSO Scheme 3 and forks/swarms (b) Residual error across different properties at two different temperatures (c) The residual error correlation between two temperatures.

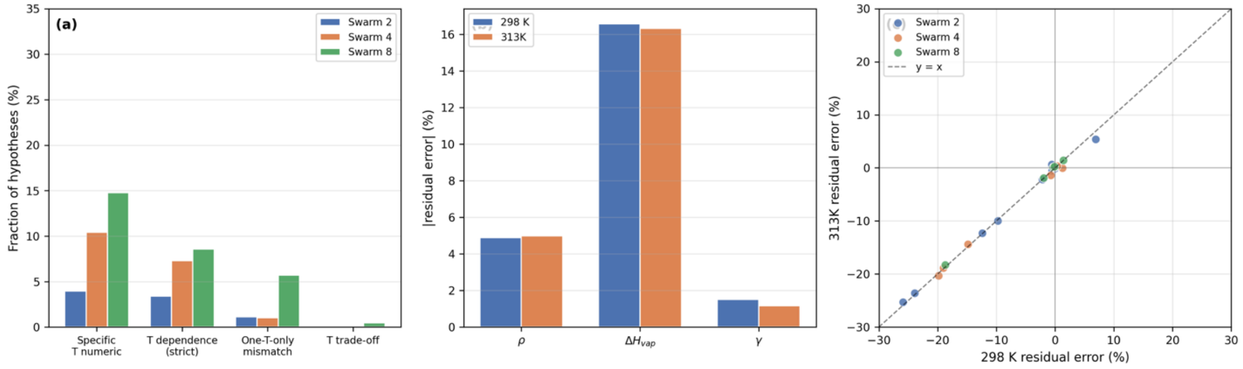


**Figure S10:** (a) Percentage of hypothesis that contained different temperature language markers across for DMA Scheme 1 and forks/swarms (b) Residual error across different properties at two different temperatures (c) The residual error correlation between two temperatures.

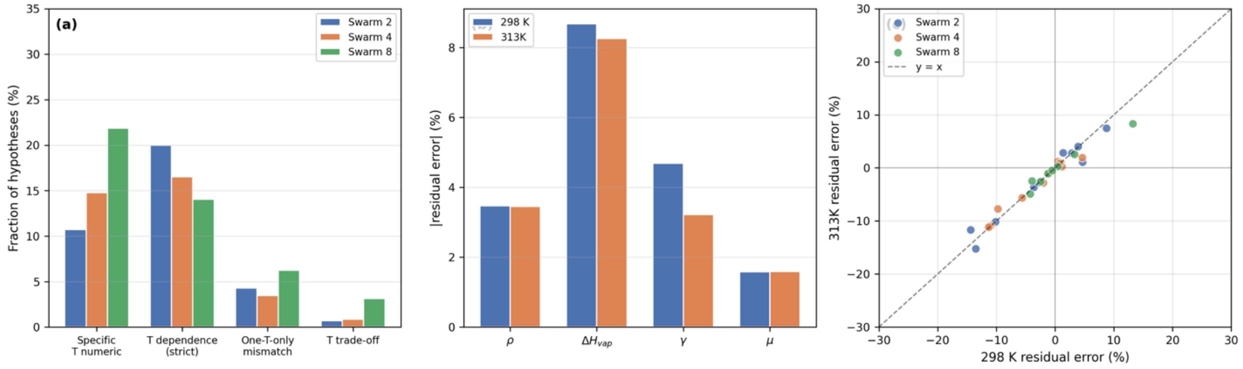


**Figure S11:** (a) Percentage of hypothesis that contained different temperature language markers across for DMA Scheme 2 and forks/swarms (b) Residual error across different properties at two different temperatures (c) The residual error correlation between two temperatures.

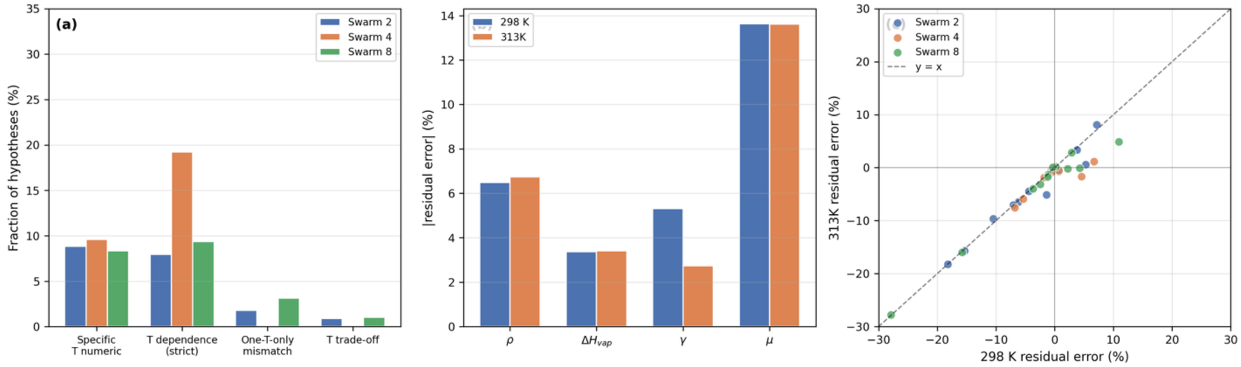


**Figure S12:** (a) Percentage of hypothesis that contained different temperature language markers across for DMA Scheme 3 and forks/swarms (b) Residual error across different properties at two different temperatures (c) The residual error correlation between two temperatures.

#### How the Hypothesis Agent Uses Multi-Fork Feedback

We scanned the “scientific rationale", “expected benefit", and “reasoning” fields of every hypothesis for four classes of language that reveal how the HA is processing prior-iteration feedback:

(a) Diagnostic Agent reference: the hypothesis text cites the DA's report directly (phrases such as 'the diagnostic reveals', 'the diagnostic report indicates', 'following the diagnostic recommendations', 'based on the diagnostic'). The DA is the framework component that aggregates all forks from the previous iteration into a single structured diagnosis, so diagnostic references are the most direct textual signature that the HA is conditioning its next move on the previous iteration's outcome.

(b) Specific numerical residual: the hypothesis cites a specific percent-error figure for a target property (phrases like '9% density deficit', 'surface tension 18% above target', 'Hvap underestimated by 12%'). Hypotheses with specific numerical residuals are conditioning on the quantitative findings of the DA rather than producing generic direction-only moves. Higher numerical-specificity rates indicate the HA is consuming richer feedback.

(c) Optimizer meta-state reference: the hypothesis references the optimizer's phase (exploration / refinement), the stuck counter, local minima, or stagnation signals. Meta-state references indicate that the HA is aware of where it is in the overall run and is modulating its reasoning accordingly.

(d) Number of sub-strategy markers in the reasoning field: a count of '(1)', '(2)', '(3)', etc. plus 'Fork 1', 'Fork 2', 'Strategy 1' markers within the HA's reasoning text. This count approximates the number of distinct exploration directions that the HA is simultaneously proposing for the OA to instantiate into concrete parameter sets. Higher values indicate that the HA is decomposing the search into more orthogonal probes per iteration.

As a negative control, we also measured

(e) backward-looking per-fork language: phrases such as 'Fork 2 showed X', 'across the forks', 'best fork', 'worst fork', 'fork variance'. If the HA were reading individual per-fork outcomes and contrasting them in its rationale, we would expect this rate to be non-trivial and to scale with fork count (since more forks give the HA more pairs to compare). The control test therefore distinguishes between the possibility that the HA reads forks directly versus the possibility that it reads a pre-aggregated DA summary. The below table shows the categorization of feedback according to the forks and solvents.

| Solvent | Fork | n hyp | DA ref. | Numeric residual | Meta-state | Backward fork (control) | Sub-strategies/hyp |
|---|---|---|---|---|---|---|---|
| **DMSO** | Fork 2 | 149 | 65.8% | 48.3% | 49.0% | 1.3% | 3.01 |
| **DMSO** | Fork 4 | 117 | 74.4% | 47.9% | 63.2% | 5.1% | 4.64 |
| **DMSO** | Fork 8 | 115 | 90.4% | 82.6% | 76.5% | 1.7% | 4.36 |
| **DMA** | Fork 2 | 113 | 79.6% | 69.9% | 63.7% | 0.9% | 3.01 |
| **DMA** | Fork 4 | 52 | 84.6% | 78.8% | 46.2% | 3.8% | 4.81 |
| **DMA** | Fork 8 | 96 | 93.8% | 63.5% | 63.5% | 2.1% | 5.47 |

The DA reference rate increases markedly with the fork count in both solvents. For DMSO it rises from 65.8% at Fork 2 to 90.4% at Fork 8, and for DMA from 79.6% to 93.8%. The HA at Fork 8 references the DA's report in more than nine out of ten hypotheses, whereas at Fork 2 it does so only about seven times in ten. At the higher fork count, the HA is much more consistently grounding its proposals in the DA's aggregated summary of the previous iteration.

The frequency with which the HA cites a specific numerical residual error follows the same monotonic pattern, rising from 48.3% to 82.6% in DMSO and 69.9% to 63.5% in DMA. Higher-fork hypotheses are more likely to explicitly quote residual errors in percentage form and to reason those errors directly, rather than producing direction-only moves ('increase ε') without numerical grounding. Combined with the DA-reference finding, this shows that the HA at higher fork counts is both reading and quoting from the DA's quantitative assessment.

The number of distinct sub-strategy markers per hypothesis scales with fork count: Fork 2 runs average 3.01 strategy markers per hypothesis in DMSO, while Fork 8 runs average 4.36. The Hypothesis Agent explicitly structures its next move as an N-way exploration when N forks are available, assigning distinct physical rationales to each slot. This is consistent with the OA downstream needing to instantiate the hypothesis into nfork concrete parameter sets, and it indicates that the HA is aware of the fork budget and uses it structurally.

Importantly, the backward-looking per-fork reference rate is very low across all fork counts. For DMSO, the rates are 1.3%, 5.1%, and 1.7% for Fork 2, 4, and 8, respectively, and for DMA, 0.9%, 3.8%, and 2.1%. The Hypothesis Agent almost never references individual per-fork outcomes in its rationale: it does not say things like 'Fork 3 showed X while Fork 5 showed Y'. Importantly, this rate does not scale with fork count: higher fork counts do not produce more per-fork comparisons in the text, even though they contain more fork pairs that could in principle be compared.

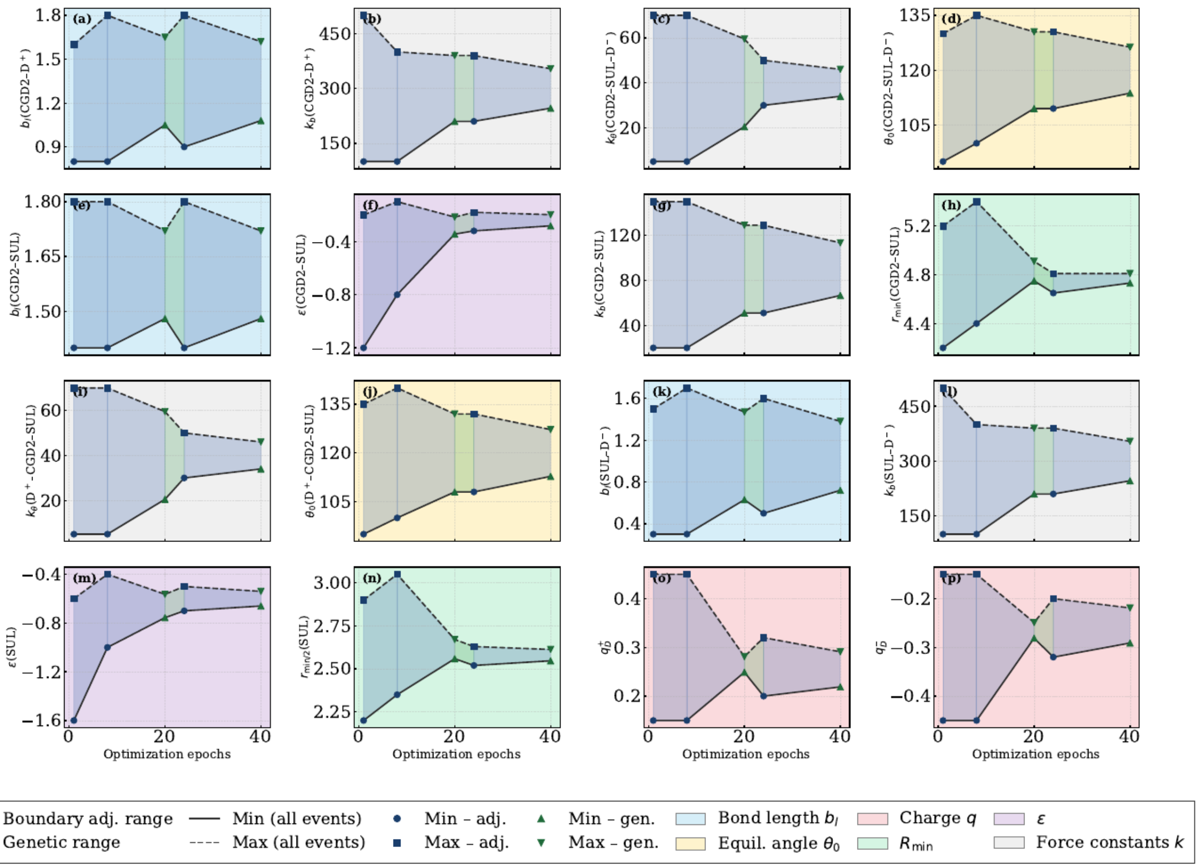


**Figure S13:** The automated parameter boundary (bounds) adjustment over the optimization epochs by the boundary agent. The green triangles denote the boundary changes due to genetic operations at a 20th-epoch interval. The progressive narrowing of the bounds reflects a genetic bottleneck phenomenon, indicating convergence toward optimal parameter combinations. The bond length, angles, charges, LJ Rmin, epsilon, and force constants boundary changes are highlighted in cyan, yellow, mauve, pink, green, and grey background colors, respectively.

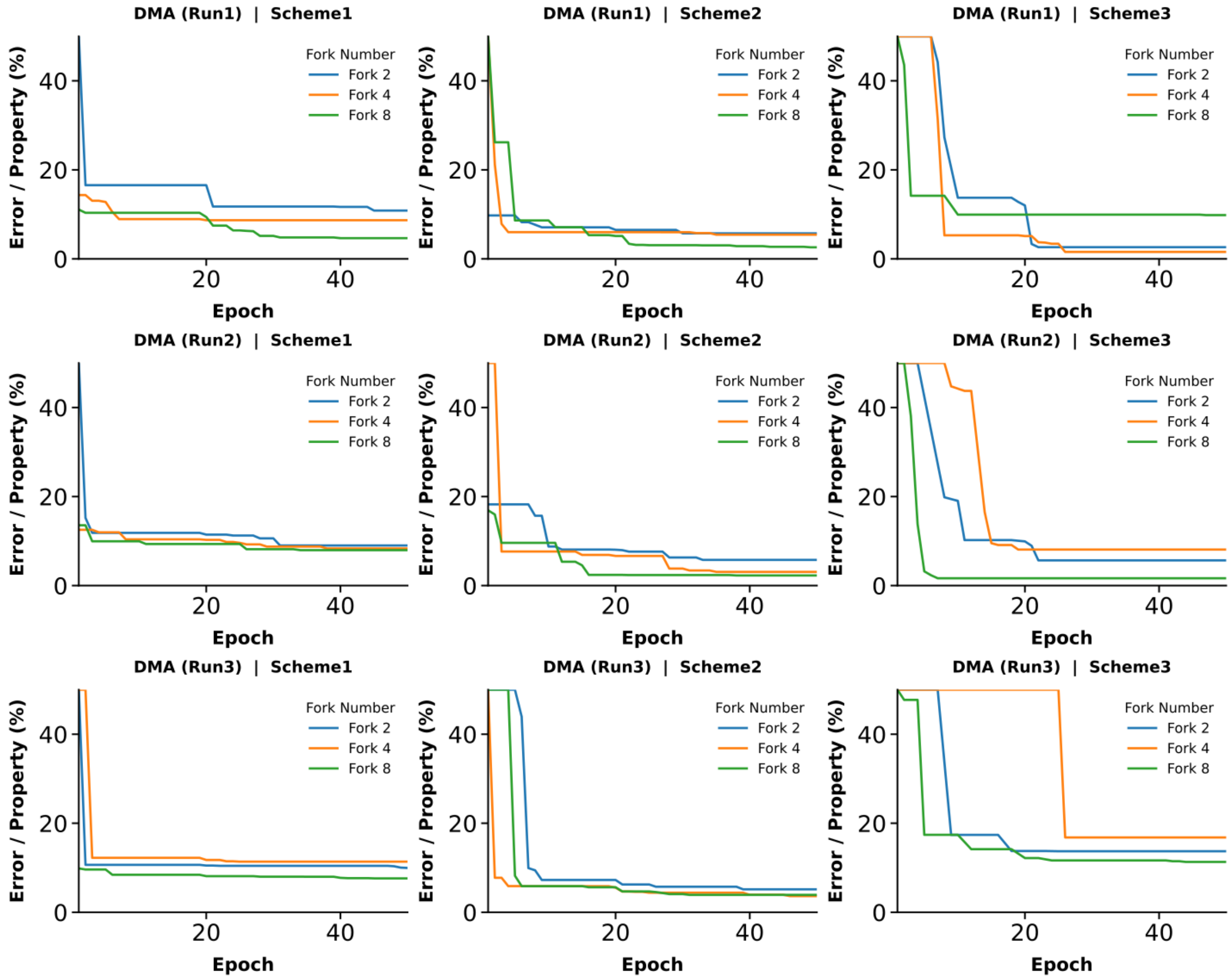


**Figure S14:** DMA percentage error per property of the global best as a function of optimization epochs for three independent runs across three mapping schemes at 298 K. The blue, orange, and green lines show the errors for forks 2, 4, and 8, respectively.

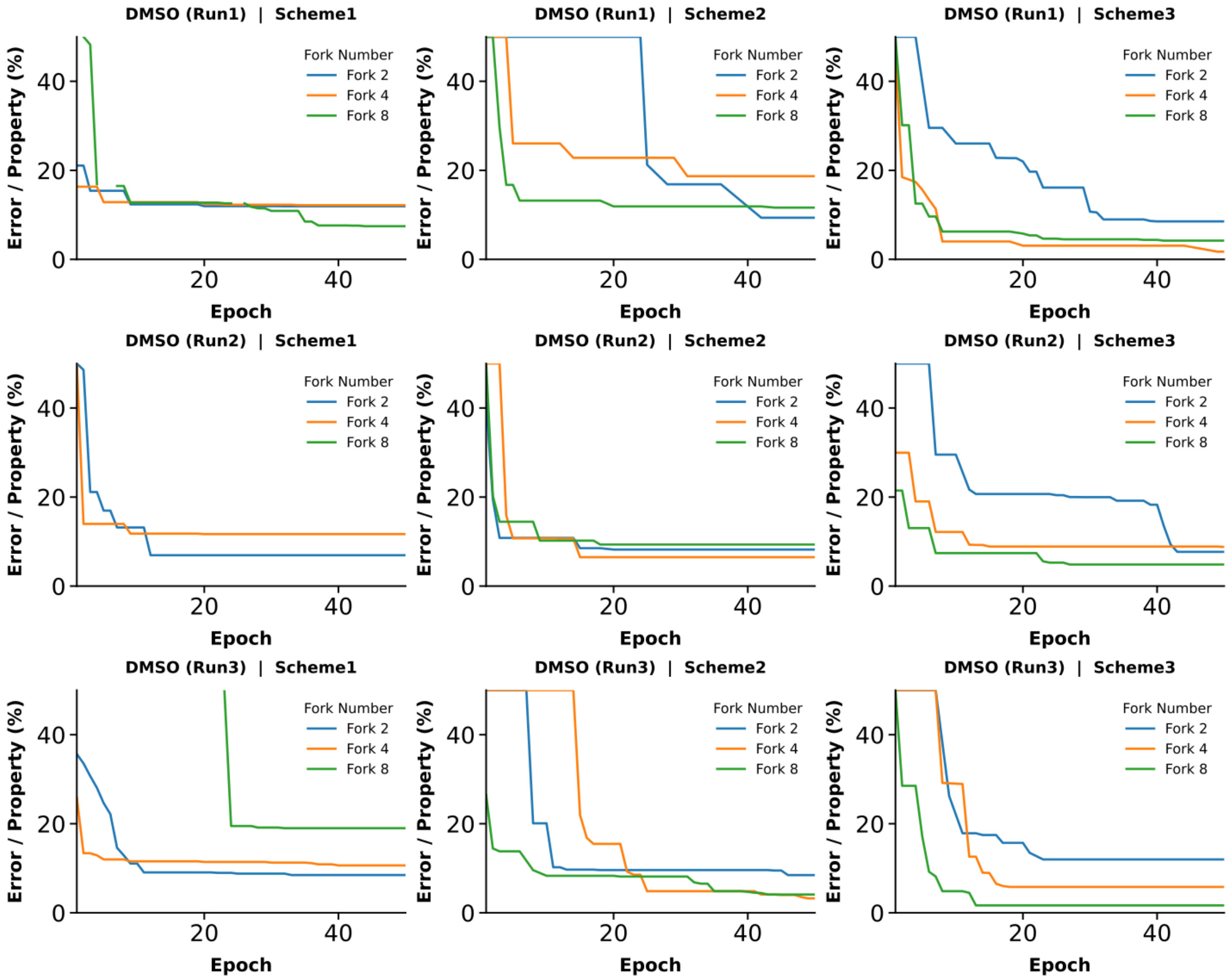


**Figure S15:** DMSO percentage error per property of the global best as a function of optimization epochs for three independent runs across three mapping schemes at 298 K. The blue, orange, and green lines show the errors for forks 2, 4, and 8, respectively.

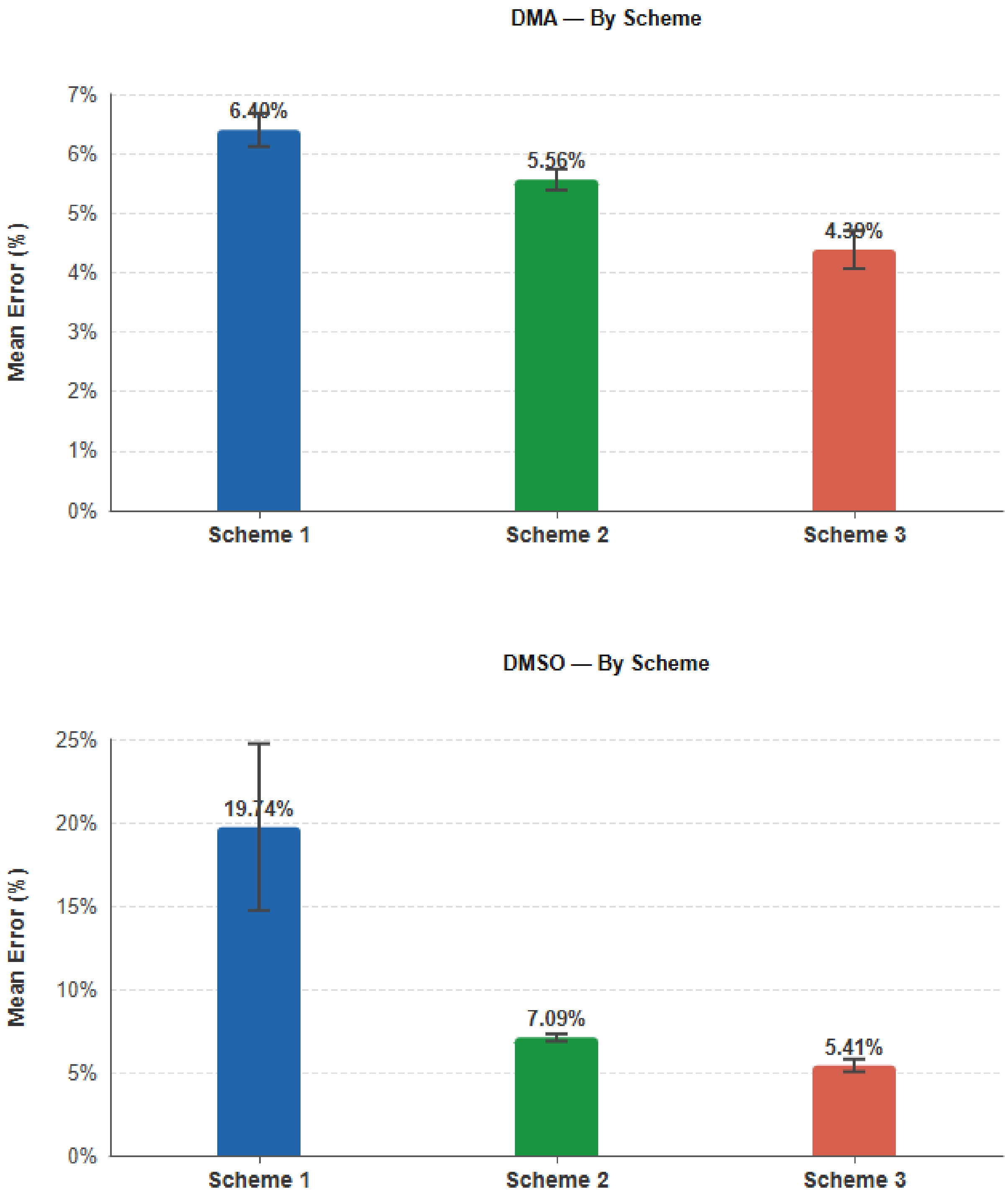


**Figure S16:** The mean error percentage of properties averaged across forks and temperatures for three different schemes for DMA and DMSO. The error bars indicate the standard deviation across three independent 100 ns of MD runs.

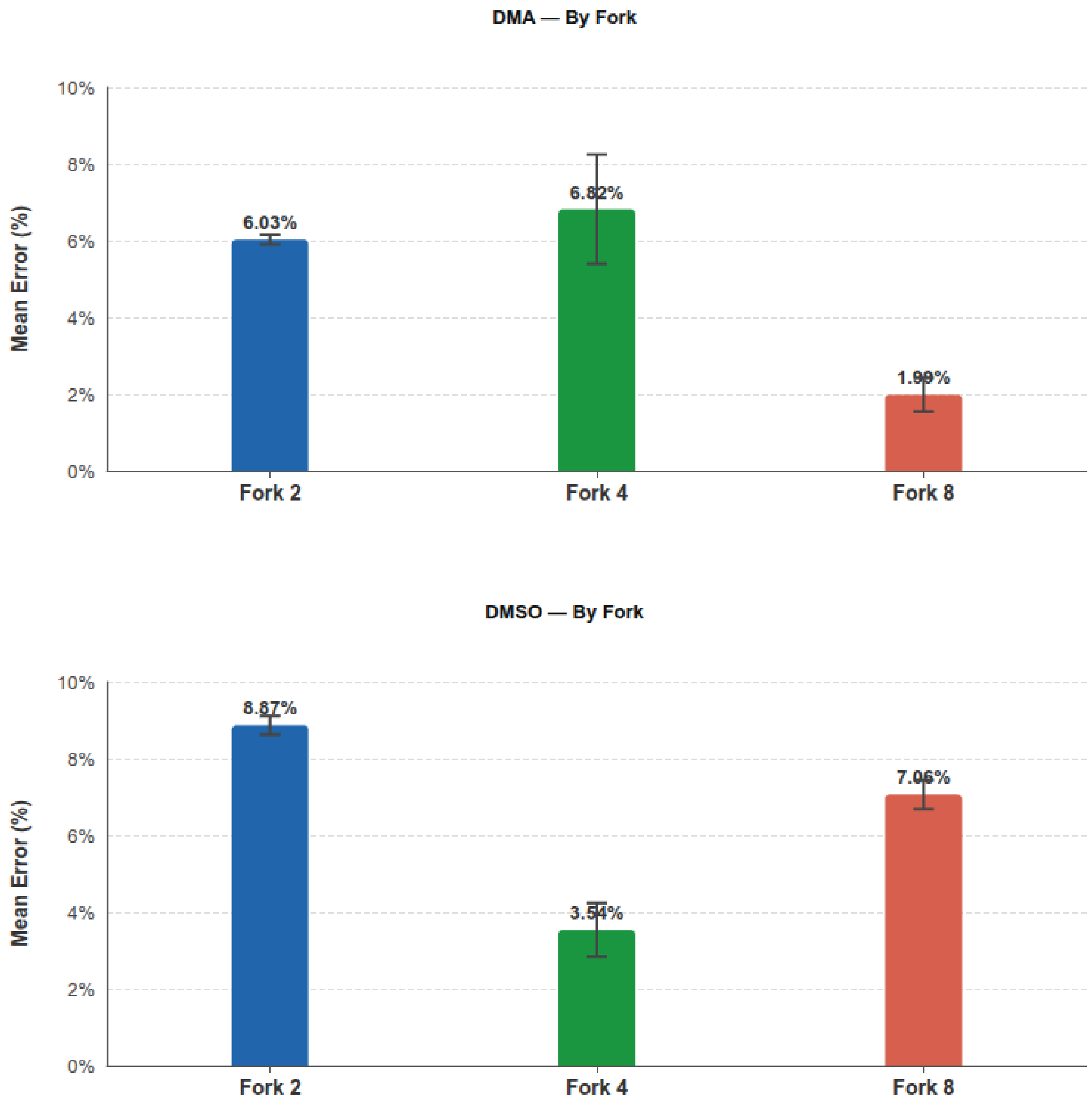


**Figure S17:** The mean error percentage of properties averaged across schemes and temperatures for three different forks for DMA and DMSO. The error bars indicate the standard deviation across three independent 100 ns of MD runs.

**Comparison of Bond and RDF among core beads:**

The bond and angle distribution of the core CG beads from three independent 100 ns MD simulations are compared against the all-atom mapped trajectory to determine the structural integrity of the CG models. The radial distribution function (RDF) is calculated within 12 Å, excluding the bonded pair peaks for clarity. For DMSO, both the bond distributions and RDFs show a close agreement for mapping Scheme 3. In contrast, for DMA mapping Scheme 3, the CGD2-CNI-MOC angle was found to be ~30% larger in the CG model compared to the all-atom reference.

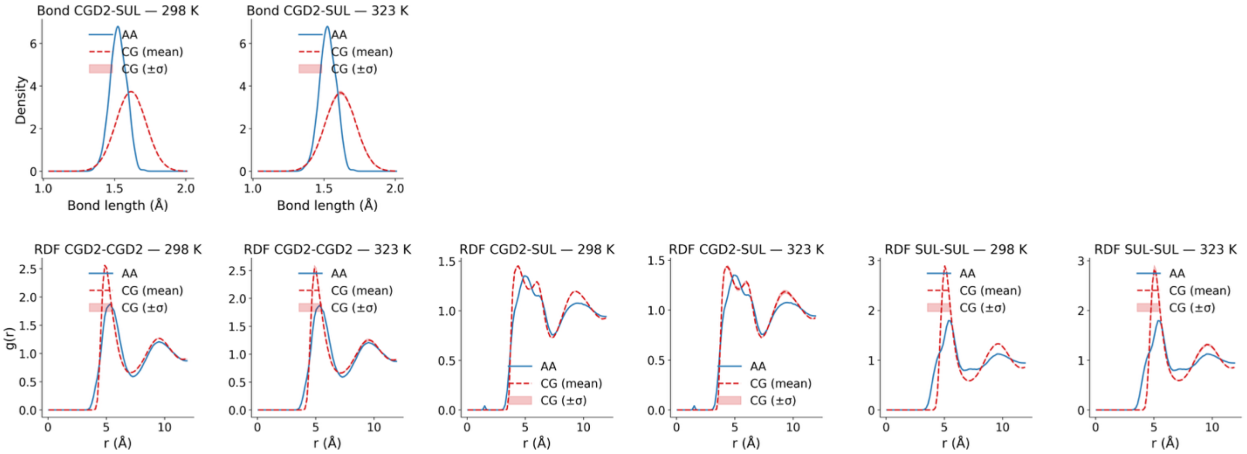


**Figure S18:** DMSO bond and RDF distributions among core beads in Scheme 3 for 298 K and 323 K are shown in red, with standard deviations in shades from three independent runs. The corresponding all-atom mapped distributions are shown in blue.

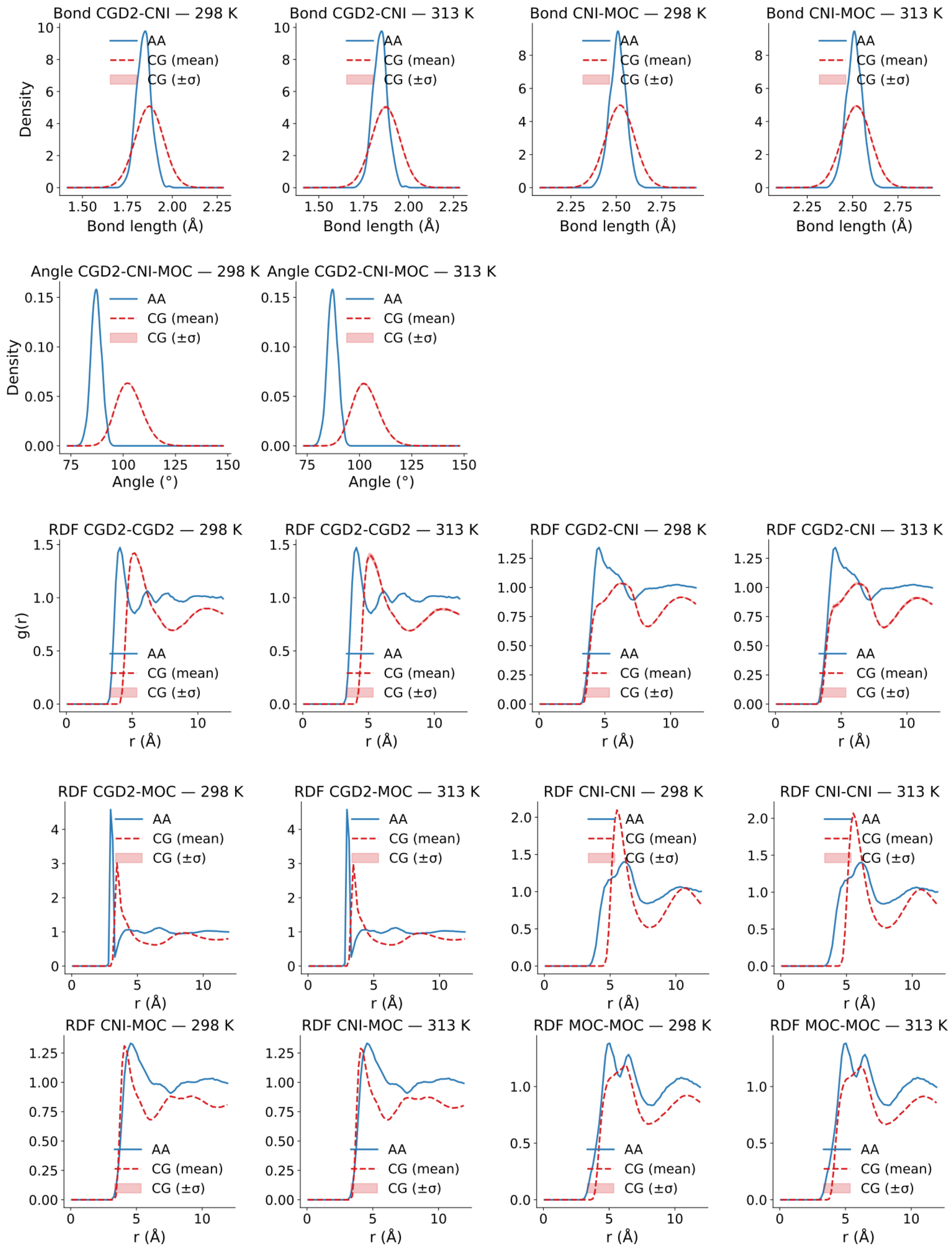


**Figure S19:** DMA bond and RDF distributions among core beads in Scheme 3 for 298 K and 313 K are shown in red, with standard deviations in shades from three independent runs. The corresponding all-atom mapped distributions are shown in blue.

**Table S2:** Tabulated thermodynamical property errors obtained from the three optimization runs for all the mapping schemes for DMA at two different temperatures, 298 K and 313 K, and DMSO at 298 K and 323 K, respectively. For each of the properties, the percentage errors are calculated with respect to the experimental targets. The green cells highlight the lowest error for each scheme.

**DMA Family**

| MAPPING SCHEME | DMA (RUN1) | | | | | DMA (RUN2) | | | | | DMA (RUN3) | | | | |
|---|---|---|---|---|---|---|---|---|---|---|---|---|---|---|---|
| | DENSITY (G CM$^{-3}$) | HEAT OF VAP. (CAL MOL$^{-1}$) | SURF. TENSION (DYN CM$^{-1}$) | DIPOLE MOM. (D) | MEAN ERROR (%) | DENSITY (G CM$^{-3}$) | HEAT OF VAP. (CAL MOL$^{-1}$) | SURF. TENSION (DYN CM$^{-1}$) | DIPOLE MOM. (D) | MEAN ERROR (%) | DENSITY (G CM$^{-3}$) | HEAT OF VAP. (CAL MOL$^{-1}$) | SURF. TENSION (DYN CM$^{-1}$) | DIPOLE MOM. (D) | MEAN ERROR (%) |
| **FORK 2** | | | | | | | | | | | | | | | |
| ***Scheme1*** | 0.54 | 25.60 | 0.61 | — | 8.92 | 2.15 | 23.77 | 0.05 | — | 8.66 | 9.85 | 12.35 | 6.15 | — | 9.45 |
| ***Scheme2*** | 2.80 | 14.37 | 2.11 | 3.63 | 5.73 | 0.81 | 10.16 | 8.11 | 3.97 | 5.77 | 2.72 | 13.02 | 2.89 | 1.23 | 4.97 |
| ***Scheme3*** | 6.32 | 0.70 | 2.93 | 0.56 | 2.63 | 3.58 | 4.41 | 7.63 | 7.04 | 5.67 | 15.54 | 10.05 | 3.27 | 18.21 | 11.77 |
| **FORK 4** | | | | | | | | | | | | | | | |
| ***Scheme1*** | 3.63 | 21.00 | 1.42 | — | 8.68 | 1.04 | 20.08 | 0.65 | — | 7.25 | 18.95 | 14.61 | 0.34 | — | 11.30 |
| ***Scheme2*** | 10.98 | 8.73 | 0.86 | 1.17 | 5.43 | 5.63 | 2.43 | 3.28 | 0.95 | 3.07 | 1.06 | 11.26 | 0.70 | 0.82 | 3.46 |
| ***Scheme3*** | 0.66 | 0.61 | 3.13 | 1.83 | 1.56 | 10.44 | 2.45 | 5.31 | 14.15 | 8.09 | 5.65 | 7.17 | 3.95 | 50.48 | 16.81 |
| **FORK 8** | | | | | | | | | | | | | | | |
| ***Scheme1*** | 2.31 | 2.43 | 4.35 | — | 3.03 | 1.93 | 18.52 | 1.41 | — | 7.29 | 0.14 | 0.24 | 0.17 | — | 0.19 |
| ***Scheme2*** | 2.52 | 4.57 | 2.92 | 0.39 | 2.60 | 5.09 | 1.73 | 0.66 | 1.60 | 2.27 | 1.15 | 3.20 | 10.77 | 0.50 | 3.91 |
| ***Scheme3*** | 3.83 | 1.22 | 7.92 | 2.86 | 3.96 | 1.46 | 2.80 | 2.17 | 0.14 | 1.64 | 15.87 | 0.22 | 1.22 | 27.84 | 11.29 |

**DMSO Family**

| MAPPING SCHEME | DMSO (RUN1) | | | | | DMSO (RUN2) | | | | | DMSO (RUN3) | | | | |
|---|---|---|---|---|---|---|---|---|---|---|---|---|---|---|---|
| | DENSITY (G CM$^{-3}$) | HEAT OF VAP. (CAL MOL$^{-1}$) | SURF. TENSION (DYN CM$^{-1}$) | DIPOLE MOM. (D) | MEAN ERROR (%) | DENSITY (G CM$^{-3}$) | HEAT OF VAP. (CAL MOL$^{-1}$) | SURF. TENSION (DYN CM$^{-1}$) | DIPOLE MOM. (D) | MEAN ERROR (%) | DENSITY (G CM$^{-3}$) | HEAT OF VAP. (CAL MOL$^{-1}$) | SURF. TENSION (DYN CM$^{-1}$) | DIPOLE MOM. (D) | MEAN ERROR (%) |
| **FORK 2** | | | | | | | | | | | | | | | |
| ***Scheme1*** | 2.37 | 31.78 | 1.53 | — | 11.89 | 2.91 | 4.99 | 12.89 | — | 6.93 | 0.24 | 22.02 | 2.56 | — | 8.27 |
| ***Scheme2*** | 17.56 | 8.55 | 1.37 | 8.33 | 8.95 | 24.76 | 1.46 | 0.99 | 0.63 | 6.96 | 8.23 | 16.24 | 8.09 | 1.26 | 8.45 |
| ***Scheme3*** | 5.56 | 6.79 | 11.58 | 6.86 | 7.70 | 5.10 | 2.61 | 19.09 | 3.97 | 7.69 | 30.73 | 3.00 | 5.99 | 8.20 | 11.98 |
| **FORK 4** | | | | | | | | | | | | | | | |
| ***Scheme1*** | 0.17 | 31.15 | 2.79 | — | 11.37 | 0.96 | 31.83 | 2.22 | — | 11.67 | 22.50 | 7.45 | 1.96 | — | 10.64 |
| ***Scheme2*** | 21.32 | 5.66 | 2.38 | 45.37 | 18.68 | 4.63 | 6.88 | 6.06 | 8.38 | 6.49 | 6.90 | 1.73 | 2.96 | 1.33 | 3.23 |
| ***Scheme3*** | 1.54 | 1.35 | 2.05 | 1.96 | 1.73 | 15.51 | 11.41 | 3.73 | 4.50 | 8.79 | 16.67 | 4.23 | 0.85 | 1.48 | 5.81 |
| **FORK 8** | | | | | | | | | | | | | | | |
| ***Scheme1*** | 2.44 | 12.02 | 4.29 | — | 6.25 | 30.81 | 2.59 | 9.90 | — | 14.43 | 0.95 | 55.24 | 0.80 | — | 19.00 |
| ***Scheme2*** | 21.06 | 7.36 | 0.61 | 16.03 | 11.26 | 11.48 | 5.70 | 15.08 | 5.07 | 9.33 | 1.91 | 10.15 | 0.38 | 2.87 | 3.83 |
| ***Scheme3*** | 7.85 | 7.59 | 1.00 | 0.37 | 4.20 | 11.79 | 0.34 | 2.73 | 4.47 | 4.83 | 1.08 | 1.29 | 4.07 | 0.16 | 1.65 |

**Table S3:** Percentage errors in properties for DMA and DMSO across three schemes and forks for the best optimization run.

| DMA | | | | | | | |
|---|---|---|---|---|---|---|---|
| **Fork** | **Scheme** | **Density (%)** | **Hvap (%)** | **Surf. Ten. (%)** | **Dipole Mom. (%)** | **Mean Err (%)** | **Best Run** |
| Fork 2 | Scheme1 | 2.15 | 23.77 | 0.05 | — | 8.6575 | Run2 |
| Fork 2 | Scheme2 | 2.72 | 13.02 | 2.89 | 1.23 | 4.9653 | Run3 |
| Fork 2 | Scheme3 | 6.32 | 0.70 | 2.93 | 0.56 | 2.6265 | Run1 |
| Fork 4 | Scheme1 | 1.04 | 20.08 | 0.65 | — | 7.2541 | Run2 |
| Fork 4 | Scheme2 | 5.63 | 2.43 | 3.28 | 0.95 | 3.0736 | Run2 |
| Fork 4 | Scheme3 | 0.66 | 0.61 | 3.13 | 1.83 | 1.5582 | Run1 |
| Fork 8 | Scheme1 | 0.14 | 0.24 | 0.17 | — | **0.1854** | Run3 |
| Fork 8 | Scheme2 | 5.09 | 1.73 | 0.66 | 1.60 | 2.2723 | Run2 |
| Fork 8 | Scheme3 | 1.46 | 2.80 | 2.17 | 0.14 | 1.6429 | Run2 |
| **DMSO** | | | | | | | |
| **Fork** | **Scheme** | **Density (%)** | **Hvap(%)** | **Surf. Ten. (%)** | **Dipole Mom. (%)** | **Mean Err (%)** | **Best Run** |
| Fork 2 | Scheme1 | 2.91 | 4.99 | 12.89 | — | 6.9306 | Run2 |
| Fork 2 | Scheme2 | 24.76 | 1.46 | 0.99 | 0.63 | 6.9595 | Run2 |
| Fork 2 | Scheme3 | 5.10 | 2.61 | 19.09 | 3.97 | 7.6933 | Run2 |
| Fork 4 | Scheme1 | 22.50 | 7.45 | 1.96 | — | 10.6386 | Run3 |
| Fork 4 | Scheme2 | 6.90 | 1.73 | 2.96 | 1.33 | 3.2316 | Run3 |

| | | | | | | | |
|---|---|---|---|---|---|---|---|
| Fork 4 | Scheme3 | 1.54 | 1.35 | 2.05 | 1.96 | 1.7260 | Run1 |
| Fork 8 | Scheme1 | 2.44 | 12.02 | 4.29 | — | 6.2499 | Run1 |
| Fork 8 | Scheme2 | 1.91 | 10.15 | 0.38 | 2.87 | 3.8284 | Run3 |
| Fork 8 | Scheme3 | 1.08 | 1.29 | 4.07 | 0.16 | **1.6485** | Run3 |

**Table S4:** Percentage property errors for DMA and DMSO across three schemes and forks for the best optimization run.

| Molecule | Scheme | Fork 2 (%) | Fork 4 (%) | Fork 8 (%) | Trend |
|---|---|---|---|---|---|
| DMA | Scheme1 | 8.66 | 7.25 | **0.19** | ↓ improves |
| DMA | Scheme2 | 4.97 | 3.07 | 2.27 | ↓ improves |
| DMA | Scheme3 | 2.63 | 1.56 | 1.64 | ↓ improves |
| DMSO | Scheme1 | 6.93 | 10.64 | 6.25 | ↓ improves |
| DMSO | Scheme2 | 6.96 | 3.23 | 3.83 | ↓ improves |
| DMSO | Scheme3 | 7.69 | 1.73 | **1.65** | ↓ improves |

**Table S5:** Dominant percentage error sources averaged over three forks for DMA and DMSO across three schemes for all the optimization runs.

| Molecule | Scheme | Worst Property | Mean (%) | 2nd Worst | Mean (%) |
|---|---|---|---|---|---|
| DMA | Scheme1 | **HoV** | **14.70** | Density | 1.11 |
| DMA | Scheme2 | **HoV** | **5.73** | Density | 4.48 |
| DMA | Scheme3 | **Density** | **2.81** | Surf. Tension | 2.74 |
| DMSO | Scheme1 | **Density** | **9.28** | HoV | 8.16 |
| DMSO | Scheme2 | **Density** | **11.19** | HoV | 4.45 |
| DMSO | Scheme3 | **Surf. Tension** | **8.40** | Density | 2.57 |

**Table S6:** Tabulated thermodynamical properties obtained from the three independent 100 ns MD runs for three different mapping schemes for DMA and two different temperatures, 298 K and 313 K, respectively. The mean and standard deviation of the three independent runs are presented. For each of the properties, the percentage errors are calculated with respect to the experimental targets and shown within the braces.

| MAPPING SCHEME | 298K | | | | | 313K | | | | |
|---|---|---|---|---|---|---|---|---|---|---|
| | DENSITY G CM⁻³ | HEAT OF VAP. KCAL MOL⁻¹ | SURF. TENSION DYN CM⁻¹ | DIPOLE D | MEAN ERR(%) | DENSITY G CM⁻³ | HEAT OF VAP. KCAL MOL⁻¹ | SURF. TENSION DYN CM⁻¹ | DIPOLE D | MEAN ERR(%) |
| *Target* | 0.9361 | 10.95 | 32.430 | 3.7200 | — | 0.9241 | 10.74 | 31.56 | 3.7200 | — |
| **Scheme1** Fork 2 | **0.9164 ± 0.0004** (2.10%) | **8.346 ± 0.021** (23.79%) | **33.402 ± 0.015** (3.00%) | **NA** | **9.63 ± 0.06 %** | **0.9122 ± 0.0071** (1.29%) | **8.294 ± 0.055** (22.80%) | **32.720 ± 0.937** (3.68%) | **NA** | **9.26 ± 0.58 %** |
| **Scheme2** Fork 2 | **0.9122 ± 0.0015** (2.56%) | **9.945 ± 0.480** (9.19%) | **35.566 ± 0.196** (9.67%) | **3.7848 ± 0.0956** (2.56%) | **5.99 ± 0.95 %** | **0.9083 ± 0.0081** (1.71%) | **9.879 ± 0.365** (8.04%) | **35.192 ± 1.608** (11.51%) | **3.7846 ± 0.0957** (2.56%) | **5.95 ± 0.68 %** |
| **Scheme3** Fork 2 | **0.8785 ± 0.0000** (6.16%) | **11.073 ± 0.091** (1.11%) | **33.094 ± 0.733** (2.42%) | **3.6990 ± 0.0022** (0.56%) | **2.56 ± 0.39 %** | **0.8735 ± 0.0082** (5.48%) | **10.975 ± 0.095** (2.16%) | **32.473 ± 0.741** (2.89%) | **3.6997 ± 0.0016** (0.55%) | **2.77 ± 0.59 %** |
| **Scheme1** Fork 4 | **0.9020 ± 0.0002** (3.65%) | **8.594 ± 0.022** (21.52%) | **33.045 ± 0.535** (1.90%) | **NA** | **9.02 ± 0.62** | **0.8909 ± 0.0002** (3.59%) | **8.464 ± 0.020** (21.21%) | **31.546 ± 0.344** (0.85%) | **NA** | **8.55 ± 0.15 %** |

| | | | | | | | | | | |
|---|---|---|---|---|---|---|---|---|---|---|
| **Scheme2** Fork 4 | **0.9016 ± 0.0002** (3.68%) | **11.045 ± 0.029** (0.86%) | **36.026 ± 0.047** (11.09%) | **3.7565 ± 0.0017** (0.98%) | **4.15 ± 0.09 %** | **0.8976 ± 0.0073** (2.87%) | **11.010 ± 0.084** (2.48%) | **35.970 ± 0.993** (13.97%) | **3.7555 ± 0.0022** (0.95%) | **5.07 ± 0.73 %** |
| **Scheme3** Fork 4 | **0.9499 ± 0.0004** (1.48%) | **11.981 ± 0.065** (9.41%) | **37.189 ± 0.802** (14.68%) | **3.9050 ± 0.0010** (4.97%) | **7.40 ± 0.79 %** | **0.9451 ± 0.0082** (2.27%) | **11.806 ± 0.100** (9.89%) | **36.830 ± 1.305** (16.70%) | **3.9047 ± 0.0005** (4.96%) | **8.46 ± 1.45 %** |
| **Scheme1** Fork 8 | **0.9349 ± 0.0001** (0.13%) | **10.833 ± 0.028** (1.08%) | **32.330 ± 0.538** (1.11%) | **NA** | **0.77 ± 0.39 %** | **0.9317 ± 0.0056** (0.82%) | **10.834 ± 0.101** (0.98%) | **31.933 ± 0.635** (1.71%) | **NA** | **1.17 ± 0.74 %** |
| **Scheme2** Fork 8 | **0.9966 ± 0.0002** (6.46%) | **12.119 ± 0.045** (10.66%) | **32.187 ± 0.767** (1.56%) | **3.8202 ± 0.0025** (2.69%) | **5.34 ± 0.36 %** | **0.9918 ± 0.0081** (7.33%) | **12.080 ± 0.043** (12.44%) | **33.179 ± 0.644** (5.13%) | **3.8156 ± 0.0007** (2.57%) | **6.87 ± 0.81 %** |
| **Scheme3** Fork 8 | **0.9123 ± 0.0002** (2.54%) | **10.341 ± 0.168** (5.57%) | **32.845 ± 0.289** (1.28%) | **3.7128 ± 0.0004** (0.19%) | **2.40 ± 0.30 %** | **0.9069 ± 0.0092** (1.86%) | **10.197 ± 0.150** (5.08%) | **32.748 ± 1.268** (3.77%) | **3.7127 ± 0.0012** (0.20%) | **2.73 ± 0.69 %** |

**Table S7:** Tabulated thermodynamical properties obtained from the three independent 100 ns MD runs for three different mapping schemes for DMSO and two different temperatures, 298 K and 323 K, respectively. The mean and standard deviation of the three independent runs are presented. For each of the properties, the percentage errors are calculated with respect to the experimental targets and shown within the braces.

| MAPPING SCHEME | 298K | | | | | 323K | | | | |
|---|---|---|---|---|---|---|---|---|---|---|
| | DENSITY G CM$^{-3}$ | HEAT OF VAP.KCAL MOL$^{-1}$ | SURF. TENSION DYN CM$^{-1}$ | DIPOLE D | MEAN ERR(%) | DENSITY G CM$^{-3}$ | HEAT OF VAP.KCAL MOL$^{-1}$ | SURF. TENSION DYN CM$^{-1}$ | DIPOLE D | MEAN ERR(%) |
| ***Target*** | **1.0950** | **12.648** | **42.090** | **3.960** | — | **1.0702** | **12.400** | **40.050** | **3.960** | — |
| **Scheme1** Fork 2 | **1.1179 ± 0.0005** (2.09%) | **13.214 ± 0.025** (4.47%) | **99.312 ± 31.979** (135.95%) | **NA** | **47.51 ± 25.35 %** | **1.1062 ± 0.0213** (3.36%) | **13.013 ± 0.342** (4.94%) | **70.302 ± 6.366** (75.54%) | **NA** | **27.95 ± 5.07 %** |
| **Scheme2** Fork 2 | **0.8492 ± 0.0000** (22.45%) | **13.361 ± 0.065** (5.64%) | **47.132 ± 0.457** (11.98%) | **3.8881 ± 0.0007** (1.82%) | **10.47 ± 0.36 %** | **0.8207 ± 0.0003** (23.32%) | **12.758 ± 0.133** (2.89%) | **43.113 ± 0.630** (7.65%) | **3.8915 ± 0.0011** (1.73%) | **8.90 ± 0.50 %** |

| | | | | | | | | | | |
|---|---|---|---|---|---|---|---|---|---|---|
| **Scheme3** Fork 2 | **1.0369 ± 0.0001** (5.31%) | **12.449 ± 0.030** (1.57%) | **51.067 ± 0.353** (21.33%) | **4.1192 ± 0.0004** (4.02%) | **8.06 ± 0.15 %** | **1.0178 ± 0.0001** (4.89%) | **12.014 ± 0.081** (3.11%) | **48.157 ± 0.384** (20.24%) | **4.1172 ± 0.0006** (3.97%) | **8.05 ± 0.40 %** |
| **Scheme1** Fork 4 | **1.0918 ± 0.0001** (0.29%) | **8.705 ± 0.012** (31.17%) | **44.921 ± 0.187** (6.73%) | **NA** | **12.73 ± 0.15 %** | **1.0845 ± 0.0124** (1.34%) | **8.641 ± 0.113** (30.31%) | **43.336 ± 2.107** (8.20%) | **NA** | **13.29 ± 1.83 %** |
| **Scheme2** Fork 4 | **1.0184 ± 0.0001** (6.99%) | **12.620 ± 0.048** (0.32%) | **46.467 ± 1.870** (10.40%) | **3.9186 ± 0.0014** (1.05%) | **4.69 ± 1.13 %** | **1.0181 ± 0.0002** (4.87%) | **12.686 ± 0.281** (2.31%) | **45.231 ± 2.156** (12.94%) | **3.9179 ± 0.0008** (1.06%) | **5.29 ± 1.55 %** |
| **Scheme3** Fork 4 | **1.0774 ± 0.0001** (1.61%) | **12.808 ± 0.128** (1.26%) | **41.678 ± 0.803** (1.61%) | **3.8824 ± 0.0018** (1.96%) | **1.61 ± 0.02 %** | **1.0693 ± 0.0133** (0.93%) | **12.700 ± 0.280** (2.52%) | **41.948 ± 1.045** (4.74%) | **3.8814 ± 0.0015** (1.99%) | **2.54 ± 1.04 %** |
| **Scheme1** Fork 8 | **1.0414 ± 0.0001** (4.89%) | **10.931 ± 0.118** (13.58%) | **45.017 ± 0.043** (6.96%) | **NA** | **8.47 ± 0.28 %** | **1.0347 ± 0.0116** (3.32%) | **10.827 ± 0.231** (12.69%) | **43.821 ± 1.606** (9.41%) | **NA** | **8.47 ± 0.67 %** |
| **Scheme2** Fork 8 | **1.0717 ± 0.0001** (2.13%) | **10.799 ± 0.062** (14.62%) | **44.589 ± 0.494** (5.94%) | **3.8330 ± 0.0006** (3.21%) | **6.47 ± 0.39 %** | **1.0633 ± 0.0149** (0.86%) | **10.681 ± 0.252** (13.86%) | **43.625 ± 1.937** (8.93%) | **3.8297 ± 0.0038** (3.29%) | **6.73 ± 0.37 %** |
| **Scheme3** Fork 8 | **1.1272 ± 0.0004** (2.94%) | **12.919 ± 0.196** (2.14%) | **48.954 ± 0.381** (16.31%) | **3.9683 ± 0.0010** (0.21%) | **5.40 ± 0.18 %** | **1.1197 ± 0.0127** (4.63%) | **12.868 ± 0.153** (3.77%) | **47.513 ± 2.126** (18.63%) | **3.9692 ± 0.0008** (0.23%) | **6.82 ± 1.91 %** |
